# Disulfide-rich, cyclic peptides from *Clitoria ternatea* protect against β-amyloid toxicity and oxidative stress in transgenic *Caenorhabditis elegans*

**DOI:** 10.1101/2021.02.07.430179

**Authors:** Neha V. Kalmankar, Hrudya Hari, Ramanathan Sowdhamini, Radhika Venkatesan

## Abstract

Neurotoxic aggregation of β-amyloid (Aβ) peptide is a hallmark of Alzheimer’s disease and increased reactive oxygen species (ROS) is known to be associated with this. Here, we report neuroprotective effects of disulfide-rich, circular peptides from *Clitoria ternatea* on Aβ-induced toxicity in transgenic *Caenorhabditis elegans*. We show that cyclotide-rich fractions from different plant tissues delay Aβ-induced paralysis in transgenic CL4176 strain expressing human muscle-specific Aβ_1–42_ gene. It also improved Aβ-induced defects in chemotaxis in CL2355 expressing Aβ_1-42_ in neuronal cells. ROS assay suggests that this is likely mediated by inhibition of Aβ oligomerization. Further, Aβ deposits were reduced in the strain, CL2006 treated with the fractions. Computational docking and molecular dynamics (MD) simulation support the findings since cyclotides bind effectively and stably to different forms of Aβ structures via hydrogen bonding and hydrophobic interactions. MD simulation further shows that cyclotides destabilize toxic amyloid assemblies. The study shows that cyclotides from *C. ternatea* could be a source of novel pharmacophore scaffold against neurodegenerative diseases.

## INTRODUCTION

Alzheimer’s disease (AD) is a chronic and progressive neurodegenerative disorder characterized primarily by extracellular plaques of beta-amyloid peptide (Aβ) deposits and neurofibrillary tangles (NFTs) in the brain ^1^. Majority of the strategies to combat AD have focused on inhibition of the amyloid aggregation, inhibition of *γ*-secretases, stabilization of the *α*-helical conformations, or elimination of the aggregated amyloid peptides ^2^. Among these strategies, identification of inhibitors of Aβ aggregates has gained prominence as the origin of neurotoxicity is often associated with the aggregation process. Despite studies investigating potential inhibitors of Aβ, only few have been approved for AD treatment, albeit considerable side-effects and only symptomatic benefits in mild-to-moderate patients ^3^. Plant metabolites such as ginsenosides from *Ginkgo biloba*, buckwheat trypsin inhibitor, cocoa peptide and others have been tested against Aβ-induced toxicity ^4-6^. These studies use transgenic *Caenorhabditis elegans* that express the human Aβ_1-42_ as a model to investigate the toxicity of Aβ. *C. elegans* is widely recognized as an excellent model organism to study AD pathology ^7-9^. Here, using *C. elegans*, we investigated the potential of cyclic peptides derived from an Indian medicinal plant in alleviating the Aβ-induced toxicity.

*Clitoria ternatea* (butterfly pea) (Fabaceae), is widely used as forage crop, natural food colorant and in traditional Ayurvedic medicine ^10,11^. Its extracts are reported to have a wide range of pharmacological activities ^12-14^. *C. ternatea* has drawn significant medicinal interest, notably for the production of “cyclotides”, a group of ultra-stable macrocyclic peptides present in all plant tissues ^15^. It is also the only known cyclotide-producing species in Fabaceae. Cyclotides are a novel class of gene-encoded, disulfide-rich, macrocyclic peptides (26-37 residues). They also form a conserved cyclic cystine knot (CCK) motif by three disulfide bonds (Cys I-Cys IV, Cys II-Cys V, Cys III-Cys VI) ^16-17^. This arrangement renders them exceptionally stable against enzymatic, chemical and thermal degradation ^18^. Peptides have few advantages over small molecules due to their low-toxicity, non-immunogenicity, better bioavailability and convenient synthesis ^19,20^. However, studies on neuroprotection by peptides against Aβ toxicity are limited ^6,21–24^.

Although cyclotides are known to be produced in several plant species, *de novo* characterization is laborious and challenging due to lower amounts, poor chromatographic resolution, difficult ionization, and deconvolution of complex structural arrangements. Nevertheless, with recent advances in RNA-sequencing technologies, peptide discovery and sequencing are relatively easier. Therefore, a combined peptidomic and transcriptomic approach is more useful and complementary in cyclotide characterization. The transcriptome of *C. ternatea* was assembled recently using RNA-seq data and 71 putative cyclotide sequences were identified ^25^. Here, we present the peptidome of *C. ternatea*, characterized by matching the masses of cyclotides obtained from MALDI-TOF experiments with the previously reported transcriptome library. *In vivo* anti-Aβ activity of cyclotides is shown using transgenic *Caenorhabditis elegans* (CL4176, CL2355 and CL2006) (Table 1) models that exhibit pathological behaviors associated with Aβ ^26^. We also performed *in silico* studies through molecular docking analyses and molecular dynamics (MD) simulations. Overall, findings from our study provides novel insights on the potential of cyclotides as Aβ inhibitors, specifically elucidating the molecular mechanisms involved in destabilization of Aβ protofibrils.

**Table 1.** Description of the transgenic *C. elegans* used in the study.

| Strains | Transgene | Expression | Phenotype |
| --- | --- | --- | --- |
| N2 | — | Control | Wild type movement |
| CL4176 | *myo-3/Aβ<sub>1-42</sub>* | Inducible in muscles | Inducible larval paralysis;<br>rapid paralysis |
| CL2006 | *unc-54/Aβ<sub>1-42</sub>* | Constitutive in muscles | Adult onset paralysis;<br>progressive paralysis |
| CL2355 | *snb-1/Aβ<sub>1-42</sub>* | Inducible in pan-neuronal | Wild type movement,<br>chemosensory deficits,<br>abnormal thrashing in<br>liquid, memory deficits |

## RESULTS

### Profiling of cyclotides from *C. ternatea*

Initial aqueous extracts were prepared from five tissues of *C. ternatea* (pods, stems, leaves, flowers and roots) and partially purified by reverse-phase C_18_ column chromatography. Each eluate was analyzed using MALDI-TOF-MS to identify cyclotide-like masses (2500-4000 Da) before further purification. Fractions eluted with 50, 70 and 100% ACN/ddH_2_O solvents showed cyclotide-like masses. These eluates were pooled (“crude extract”) for each plant tissue and further purified by preparative RP-HPLC (Figure 1). Early (hydrophilic) and late (hydrophobic) eluting fractions were manually collected and separated into five fractions (A-E). Each fraction was analyzed for cyclotides via MALDI-TOF MS using characteristic m/z values (2500-4000 Da). Fraction D was found to be richer in cyclotides compared to fractions A, B, C and E (Figure 1, Figure S1, Supporting Information). Fraction D for each tissue is henceforth termed as cyclotide-rich fraction (CRF) and used in all the subsequent assays. The MALDI-TOF spectra for the CRFs from five tissues of *C. ternatea* are shown in Figure 2. Using the transcriptome assembly of *C. ternatea* ^25^, we identified various cyclotide sequences by correlating the calculated mass of cyclotide genes and experimentally identified monoisotopic masses from MS data (Figure 2; Table S1, Supporting Information). Out of the 67 cyclotide-like masses identified from all five tissue CRFs, 22 of them could be sequenced using nucleotide information (Table 2).

**Table 2.**
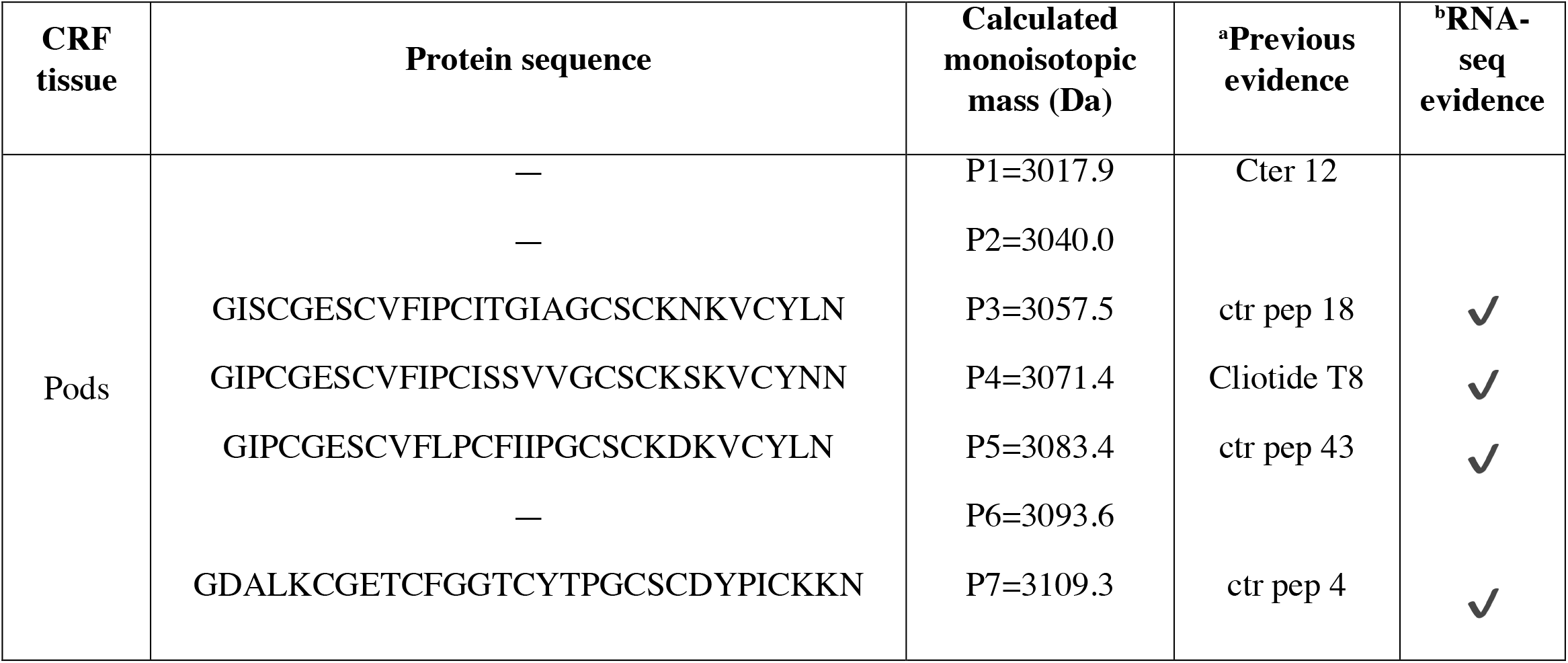
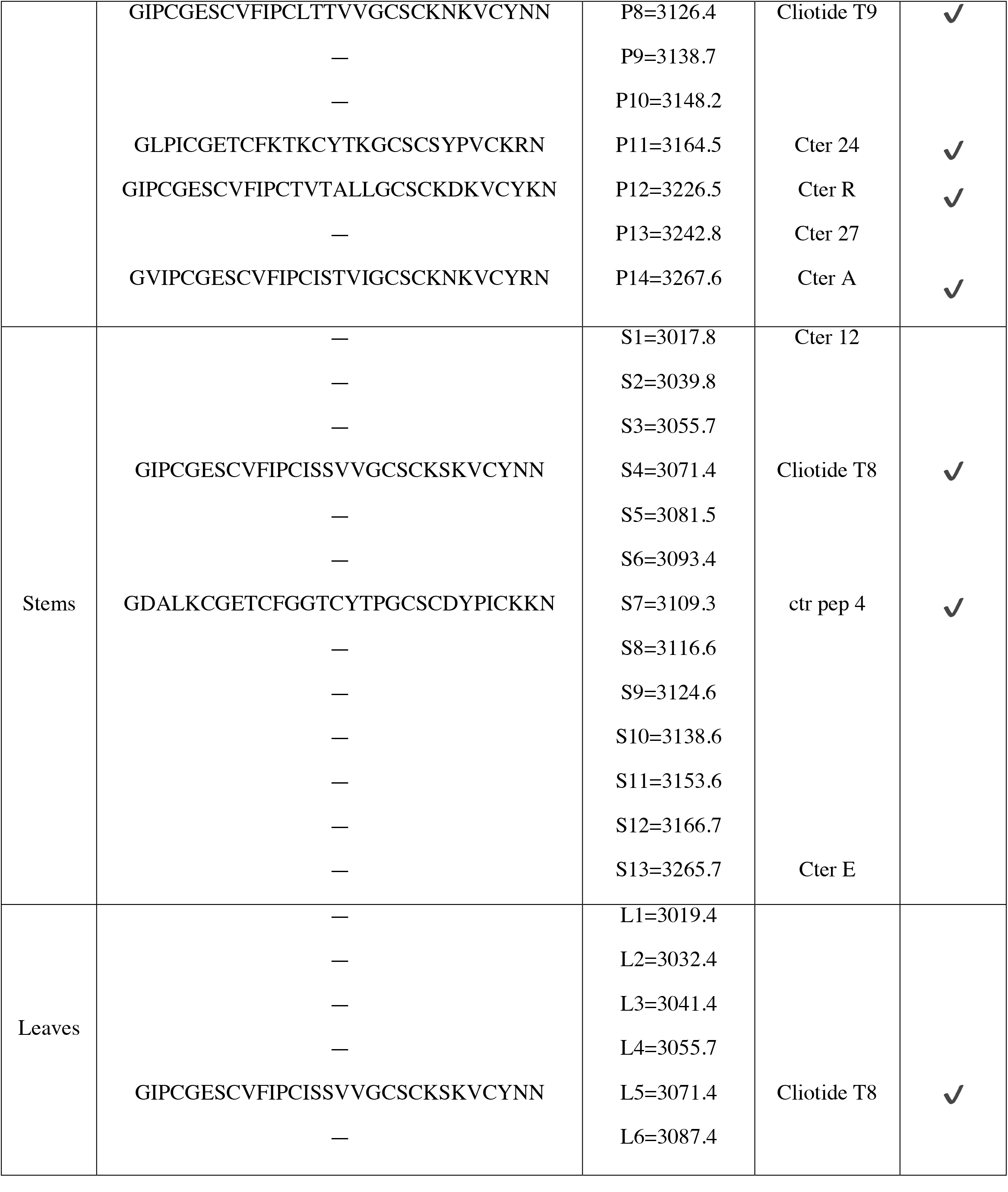
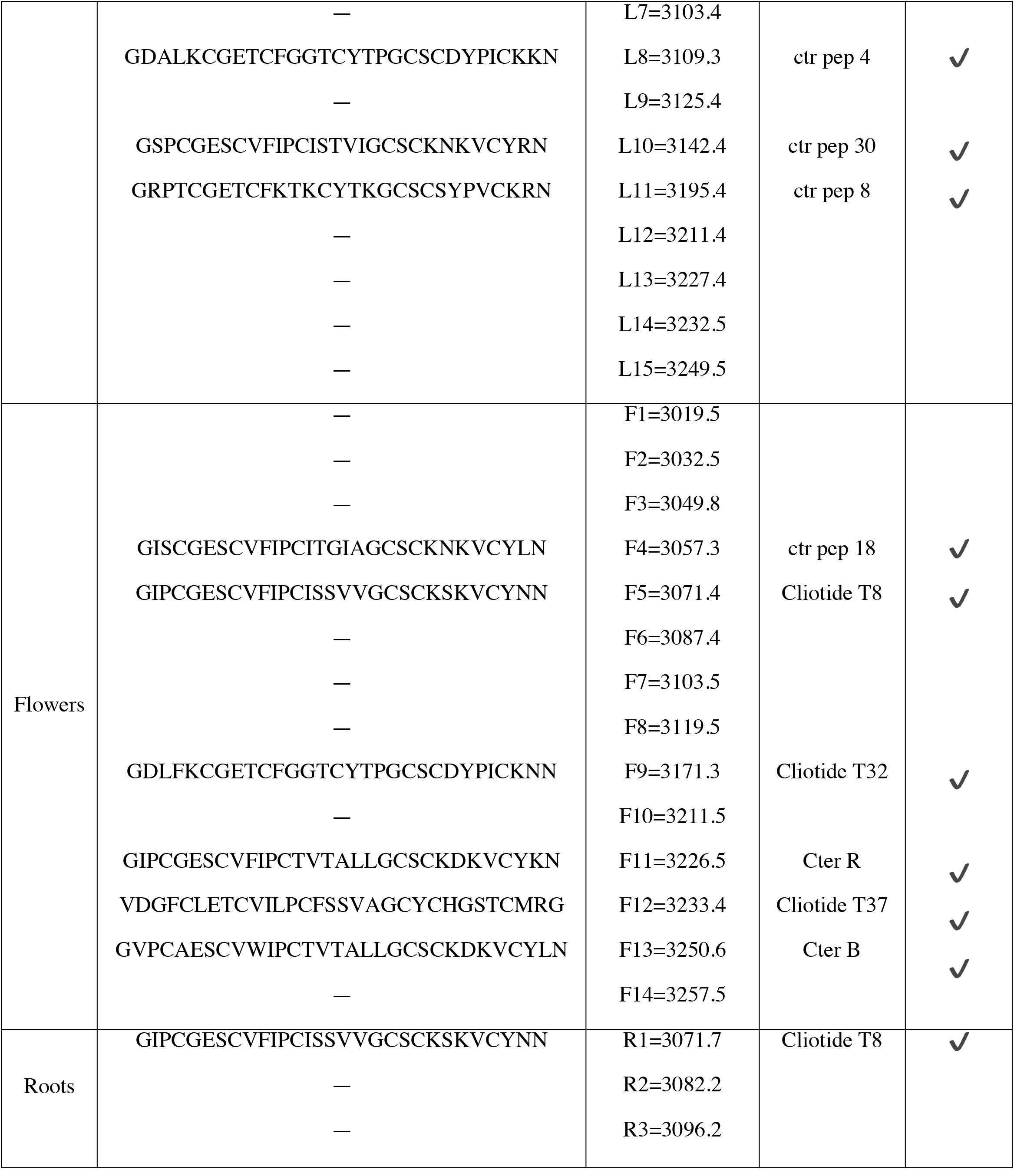
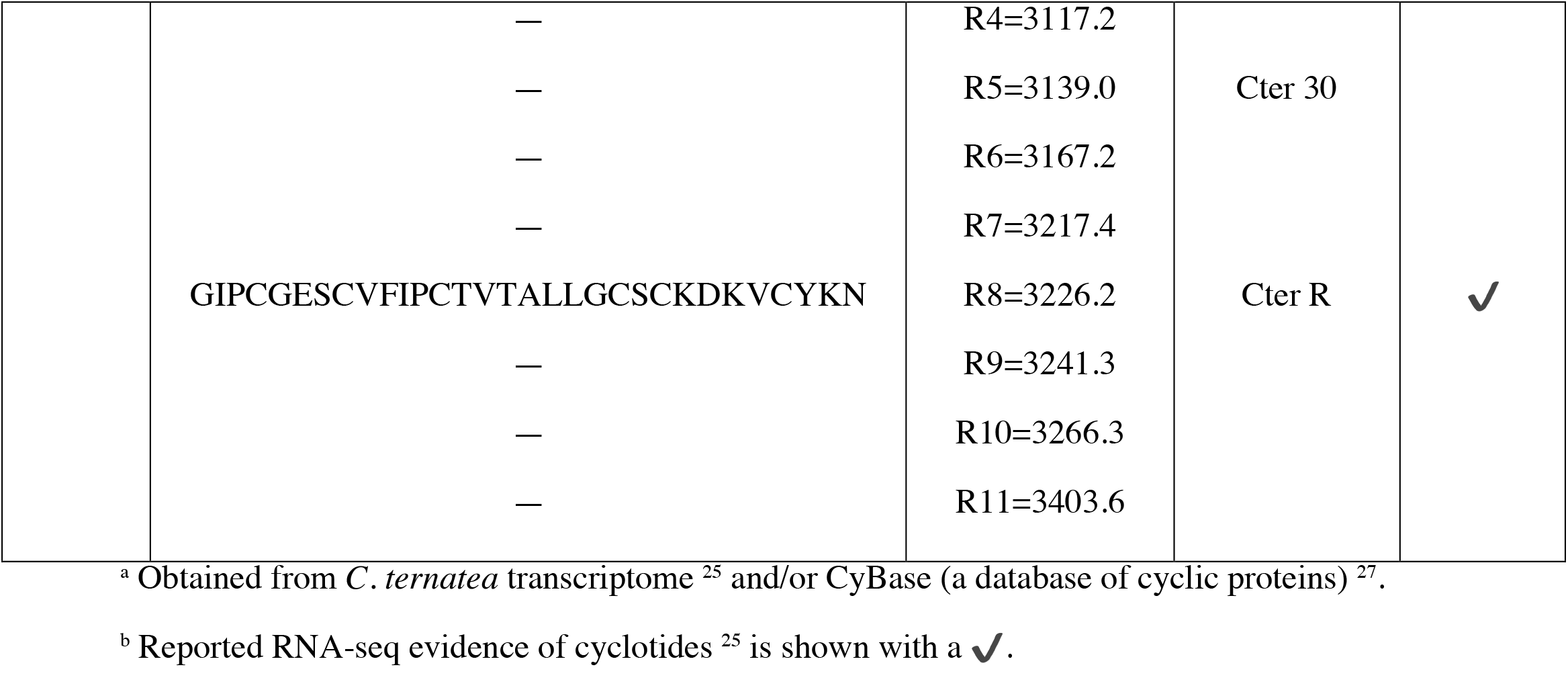
*C. ternatea* cyclotide sequences determined by proteomic and transcriptomic mining

**Figure 1.**
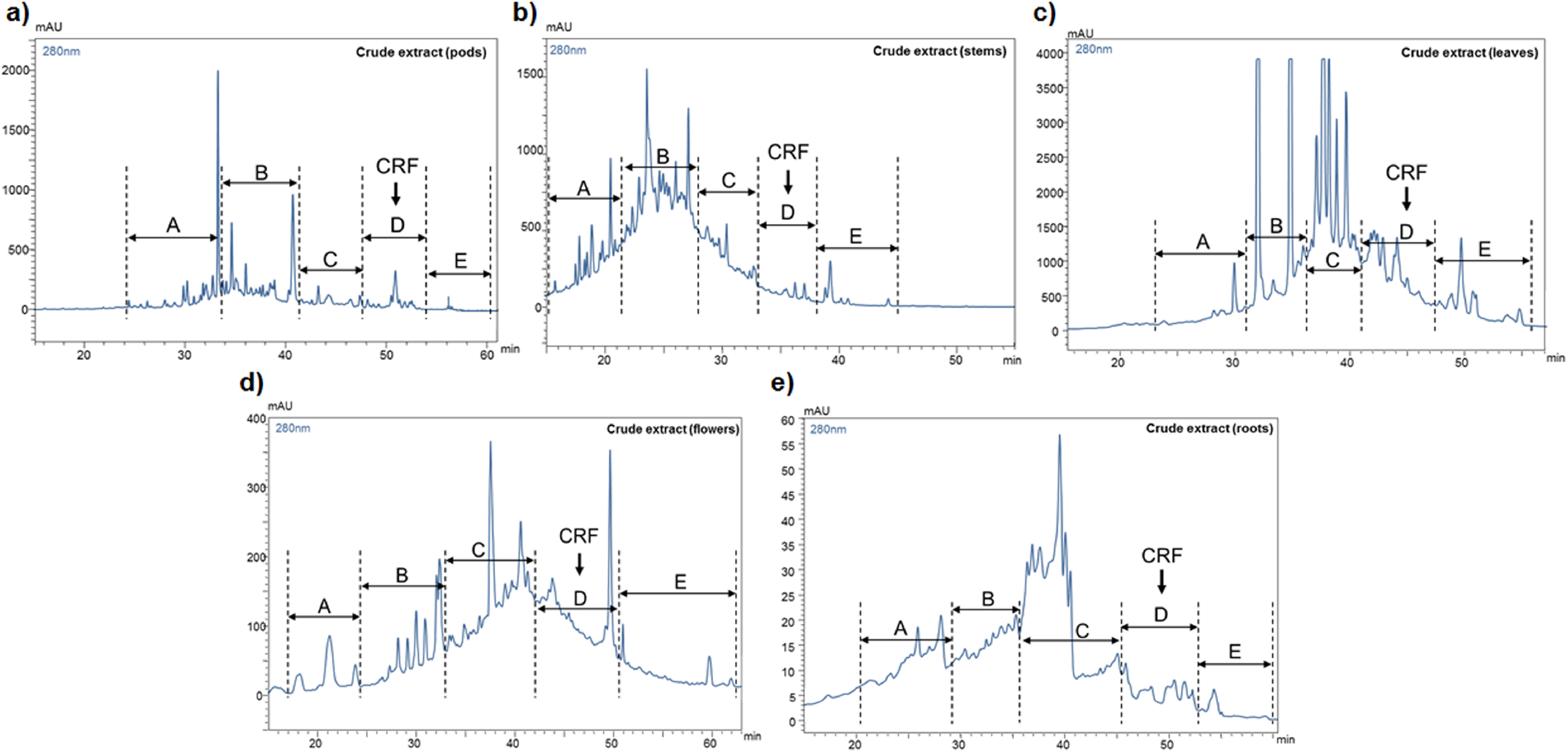
Semi-prep HPLC profiles of crude cyclotide extracts obtained from (a) pods, (b) stems, (c) leaves, (d) flowers and (e) roots of *C. ternatea*. A linear gradient of 1% min^-1^ of 0–95% buffer B (100% acetonitrile, 0.1% trifluoroacetic acid) was applied, and the eluents were monitored at 220, 254, and 280 nm. Five fractions (A-E) were collected from each plant tissue sample. The cyclotide-rich fraction (CRF or fraction D) is highlighted with an arrow.

**Figure 2.**
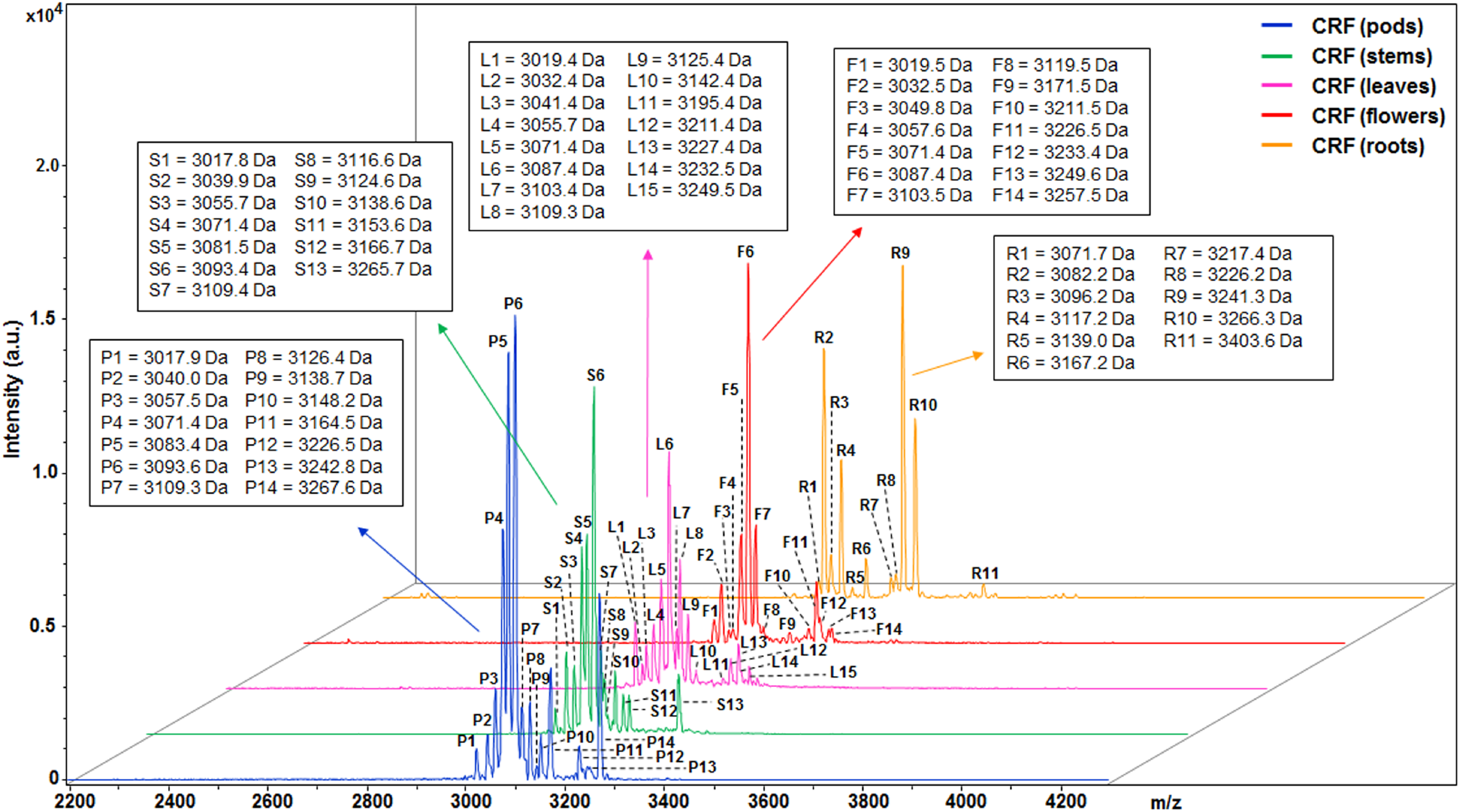
A stacked plot of MALDI-TOF mass-spectra of cyclotide-rich fraction (CRF) from different tissues of *C. ternatea*. The monoisotopic signals (M^+^= m/z - H^+^) are labeled with names and their corresponding molecular weights are highlighted in the insets.

### Effect of cyclotides on Aβ-induced paralysis in transgenic AD *C. elegans*

CRF from different tissues of *C. ternatea* were characterized for their effect on Aβ-induced paralysis in *C. elegans* strain CL4176. This strain expresses full length Aβ_1-42_ protein in the body wall muscles and shows a phenotype of Aβ expression and aggregation in muscle upon temperature upshift from 15°C to 23°C, leading to progressive paralysis. Figure 3a illustrates the duration of treatment and paralysis assays in CL4176 and wild type-strain N2. Figure 3b shows microscope images of CL4176 worms when normal and when paralyzed. Our results show that in untreated control, the worms show complete paralysis within 100 h of temperature upshift. By comparison, the paralysis rate induced by Aβ was delayed in CL4176 worms treated with CRF by ∼40 % (Figure 3c-g), irrespective of which plant tissue it was extracted from. Based on previous reports, Vitamin C was chosen as a positive control for all the assays ^28-30^. 80% of the untreated worms became paralyzed after 80 h, while only 50% of the Vitamin C treated worms were paralyzed. CRF from all the five tissues delayed the onset of paralysis in a dose–response manner. The highest concentration of CRF, i.e. 200 μg/ml concentration displayed significant delay in paralysis compared to the lower concentrations of cyclotides tested (Figure 3h). For example, when worms were treated with 200 μg/ml CRF from pods, ∼56% of the worms were paralyzed at 84^th^ hour post temperature upshift, whereas, ∼47% and ∼27% of the worms were paralyzed when treated with lower concentrations (80 μg/ml and 20 μg/ml) CRFs. This trend was observed in all tissue wise treatments, where 200 μg/ml concentration of CRF showed maximum desired activity (Figure 3h). As shown in Figure 3h, among CRF from different tissues, treatment with pods CRF (concentration: 200 μg/ml) showed a significant delay in paralysis rates, (∼ 56%) compared to the positive control (∼46%) (p = 0.041; n = 10) (Table S2, Supporting Information). These observations suggest that cyclotides have the potential to protect against Aβ-induced toxicity due to their ability to suppress paralysis in the whole animal. The wild type N2 strain showed no paralysis at all at any time, regardless of treatment (Figure 3c-g).

**Figure 3.**
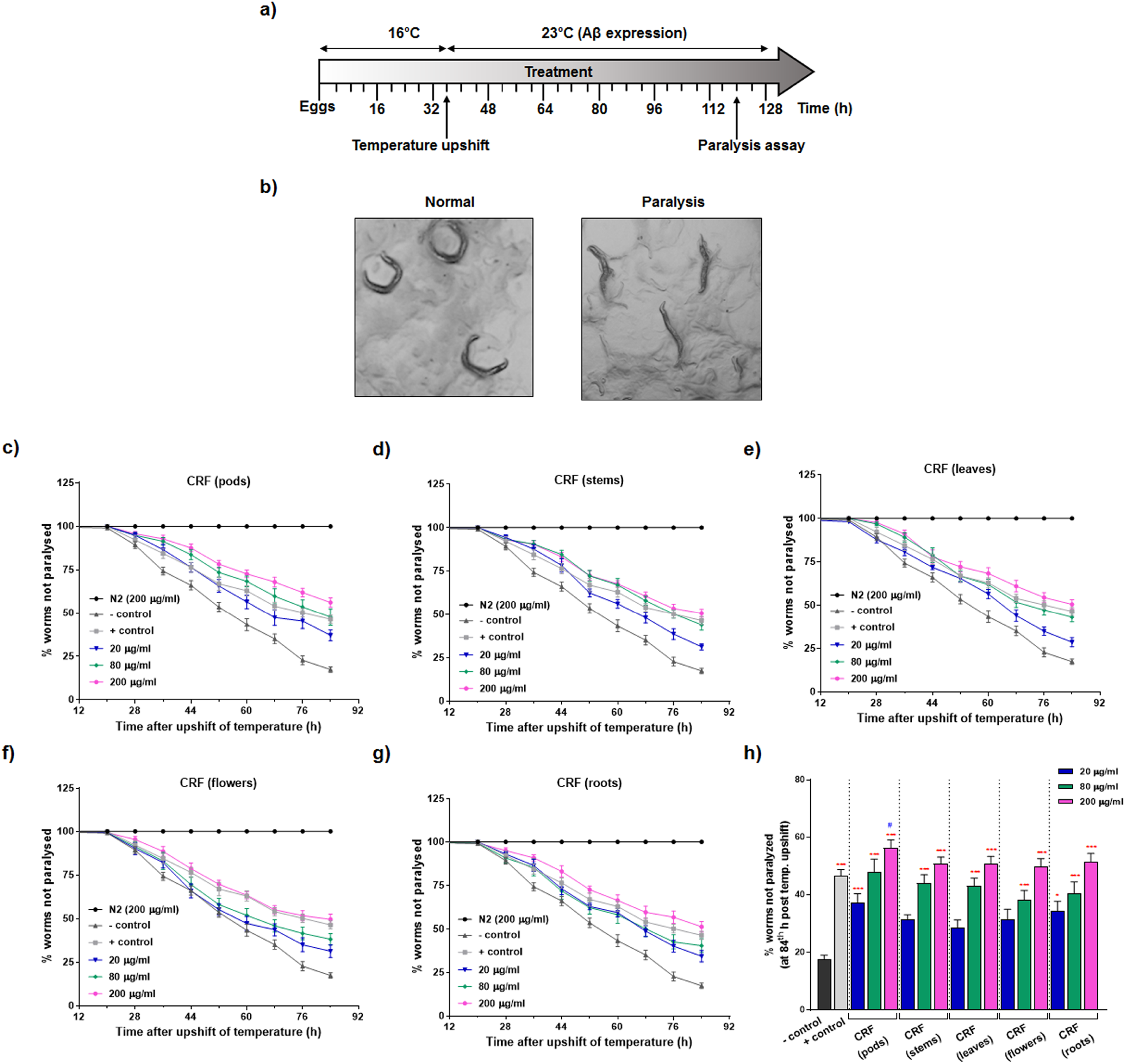
Effect of CRF from different tissues of *C. ternatea* on Aβ-induced paralysis in *C. elegans* strain CL4176 and wild type strain N2. (a) An illustrative diagram indicating the duration of treatment and paralysis assays in CL4176 and wild type strain N2 worms. (b) Microscope image showing CL4176 worms when normal and paralyzed. (c-g) Time course of Aβ-induced paralysis in the transgenic CL4176 strain and wild type N2 strain treated with a vehicle (negative control) or CRFs (20, 80, 200 μg/ml) from different tissues or Vitamin C (positive control). (h) The rate of paralysis at 84^th^ hour post temperature upshift for the treated and untreated transgenic worms. Data represent mean ± SE % paralysis from at least 10 independent experiments. Red and blue symbols represent statistically significant differences between CRF treatment versus negative and positive control groups respectively (p < 0.05: ^*^/^#^, p < 0.01: ^**^/^##^, p < 0.001: ^***^/^###^).

### Effect of cyclotides on Aβ-induced chemotaxis defects in transgenic AD *C. elegans*

In AD, Aβ accumulation and aggregation is essentially toxic to the neuronal cells and muscle specific Aβ expressing *C. elegans* model cannot specifically reflect the pathology of AD in the brain. Therefore, we assessed the neuroprotective effects of cyclotides on Aβ-induced toxicity in the neurons of *C. elegans*. We used transgenic strain CL2355 that has an inducible, pan-neuronal expression of Aβ_1-42_, to investigate whether cyclotides can rescue the chemosensory deficits caused by Aβ-induced toxicity. Details of the experiment are illustrated in Figure 4a. Using chemotaxis index (CI) as a measure to evaluate attractive (positive value) and repulsive (negative value) response, we tested the CRF from different tissues of *C. ternatea* (Figure 4b-f). CRFs from all the tissues showed significant protection of neuronal cells against amyloid toxicity compared to untreated nematodes (negative control) (Figure 4b-f). Vitamin C which was used as a positive control in the experiment showed the maximum reversal of chemotaxis defects. Amongst CRF treatment, most significant rescue was observed when CL2355 worms were treated with 200 μg/ml leaves CRF (CI positive control = 0.27 ± 0.13; CI leaves CRF = 0.26 ± 0.15) (Table S3, Supporting Information). Additionally, the effect of CRFs on the CI was dose-dependent. 200 μg/ml concentration displays significant rescue in all except in the stems CRF treatment (Figure 4c). These results suggest that cyclotides could protect neuronal cells against Aβ induced toxicity, thereby rescuing the associated defects in chemotaxis behavior.

**Figure 4.**
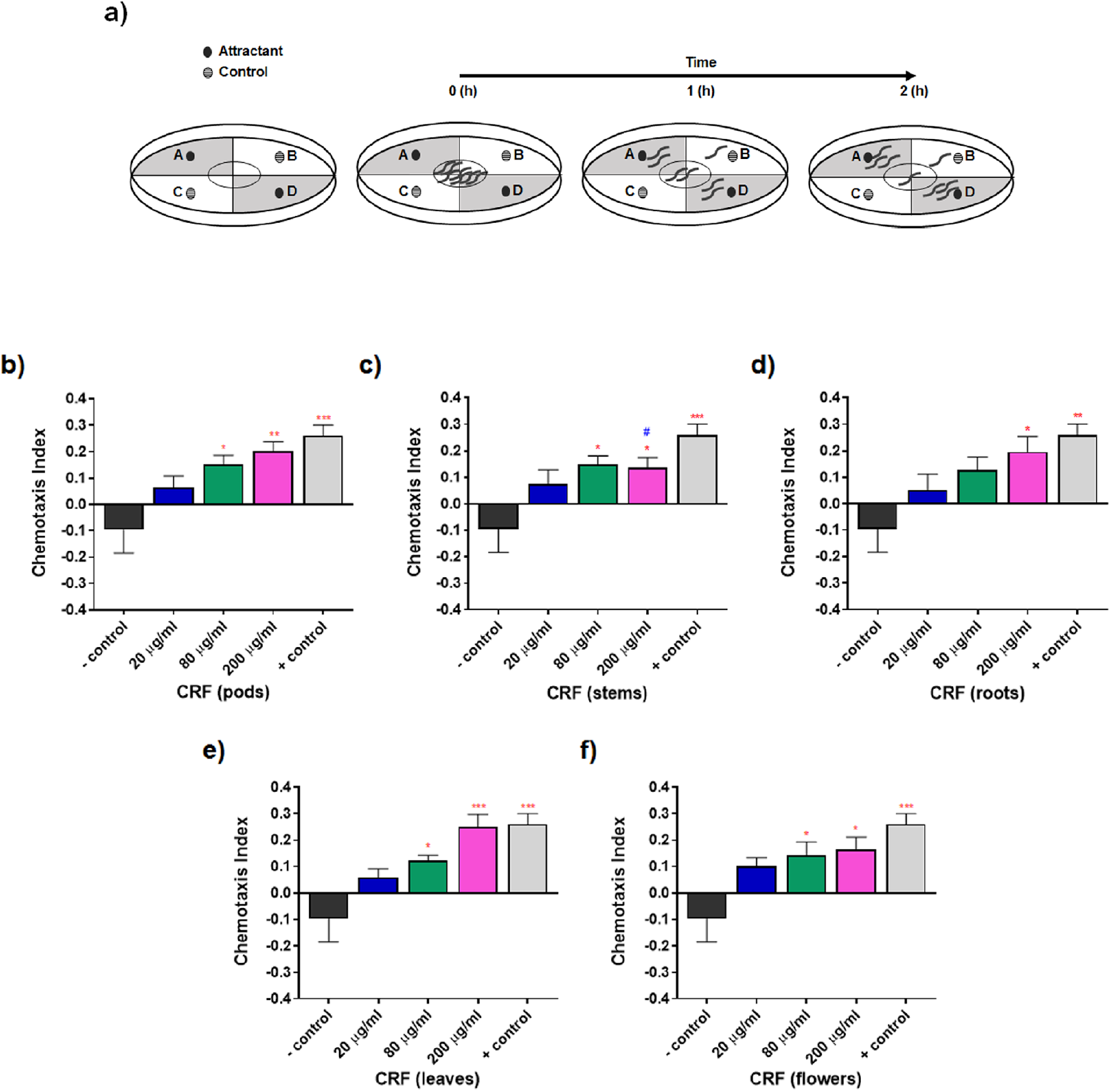
Effect of -CRF from different tissues of *C. ternatea* on Aβ-induced defect in odorant preference in *C. elegans* strain CL2355 and wild type strain N2. (a) Experimental design for the chemotaxis behavior assay performed with transgenic *C. elegans* strain CL2355 (neuronal Aβ strain) and wild type strain N2 using 0.1% benzaldehyde in 100% ethanol as an attractant and 100% ethanol as a control odorant. (b-f) Chemotaxis behavior assay on transgenic *C. elegans* strain CL2355. Data represent mean ± SE of CI from at least 10 independent experiments. Red and blue symbols represent statistically significant differences between CRF treatment versus negative and positive control groups respectively (p < 0.05: ^*^/^#^, p < 0.01: ^**^/^##^, p < 0.001: ^***^/^###^).

### Effect of cyclotides on ROS in transgenic AD *C. elegans*

From the above results, dose-responsive protective effects of CRF on Aβ-induced toxicity is evident. Next, to evaluate the antioxidant effect of cyclotides in vivo, intracellular ROS levels were measured after feeding transgenic strain CL4176 with 200 μg/ml concentration of CRFs from different tissues. Significantly lower fluorescence intensities were observed in nematodes treated with CRFs compared to untreated control nematodes expressing Aβ (Figure 5a-c) (Table S4; Supporting Information). Specifically, treatment with 200 μg/ml of CRF from flowers of *C. ternatea* exhibited the most significant reduction of intracellular ROS (∼20% reduction from untreated worms) (Figure 5c; Table S4, Supporting Information). This reduction was higher (∼7%) than positive control as well. These results suggest indirectly that the delay observed in Aβ induced paralysis of CL4176 nematodes after treatment with CRF could be due to reduced accumulation of intracellular ROS. These results establish that cyclotides possibly have antioxidant properties.

**Figure 5.**
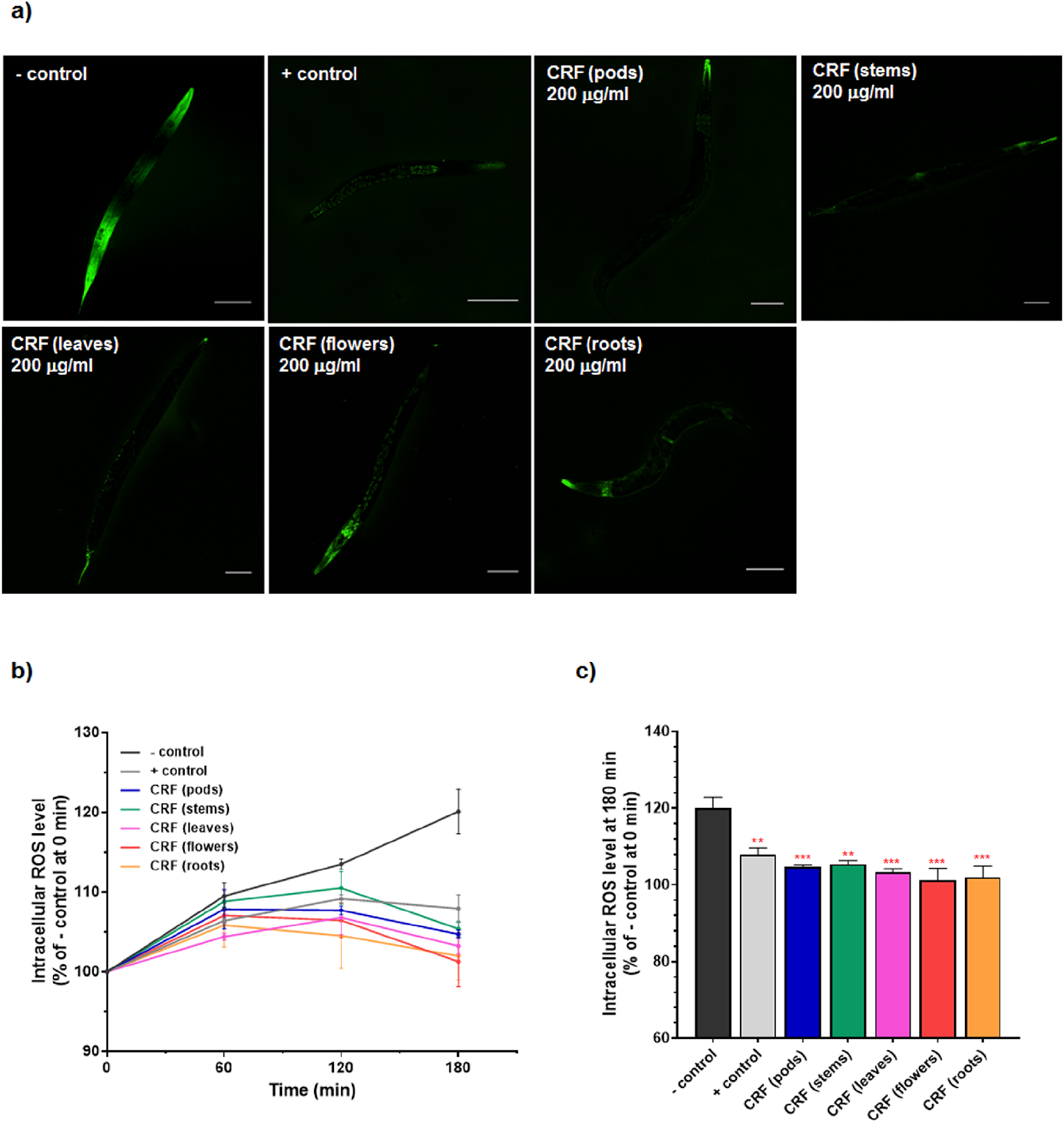
Effect of cyclotides on the intracellular reactive oxygen species (ROS) production in transgenic *C. elegans* CL4176 strain. Age-synchronized worms were treated with or without CRF, followed by the DCF assay for ROS estimation. (a) DCF fluorescent images of nematodes untreated (negative control) versus treatment with CRF from different tissues of *C. ternatea*. Scale bars: 100 μm. (b) Time-course of increase in DCF fluorescence detected by a microplate reader at 485 nm excitation and 530 nm emission. Results are expressed as percentage of fluorescence (intracellular ROS level) relative to untreated control (negative control) at t=0 which is set as 100%. (c) Corresponding % DCF fluorescence observed at t=180 min. Data represent mean ± SE of % DCF fluorescence from 4 replicates of experiments. Red and blue fonts represent statistically significant differences between CRF treatment versus negative and positive control groups respectively (p < 0.05: ^*^/^#^, p < 0.01: ^**^/^##^, p < 0.001: ^***^/^###^).

### Effect of cyclotides on Aβ deposits in transgenic AD *C. elegans*

To investigate the effect of cyclotides on Aβ aggregation, we performed thioflavin-S (ThS) fluorescence assay to visualize the formation of amyloid fibrils. Transgenic *C. elegans* strain CL2006, which expresses an Aβ protein fragment involved in the development of AD, was used for this assay. The strain shows a phenotype of constitutive Aβ expression and aggregation in muscles, leading to progressive paralysis. CL2006 worms were subjected to Aβ staining using thioflavin-S at the end of CRF treatment. Fluorescent images of the head region of the CL2006 worms shows significant reduction in Aβ deposits (blue arrowheads) in worms treated with CRFs (concentration: 200 μg/ml) compared to untreated worms (negative control) (Figure 6). The wild type N2 strain shows no Aβ deposition in the whole animal. These results suggest that cyclotides inhibit Aβ aggregation and deposition in the transgenic worms’ muscle cells.

**Figure 6.**
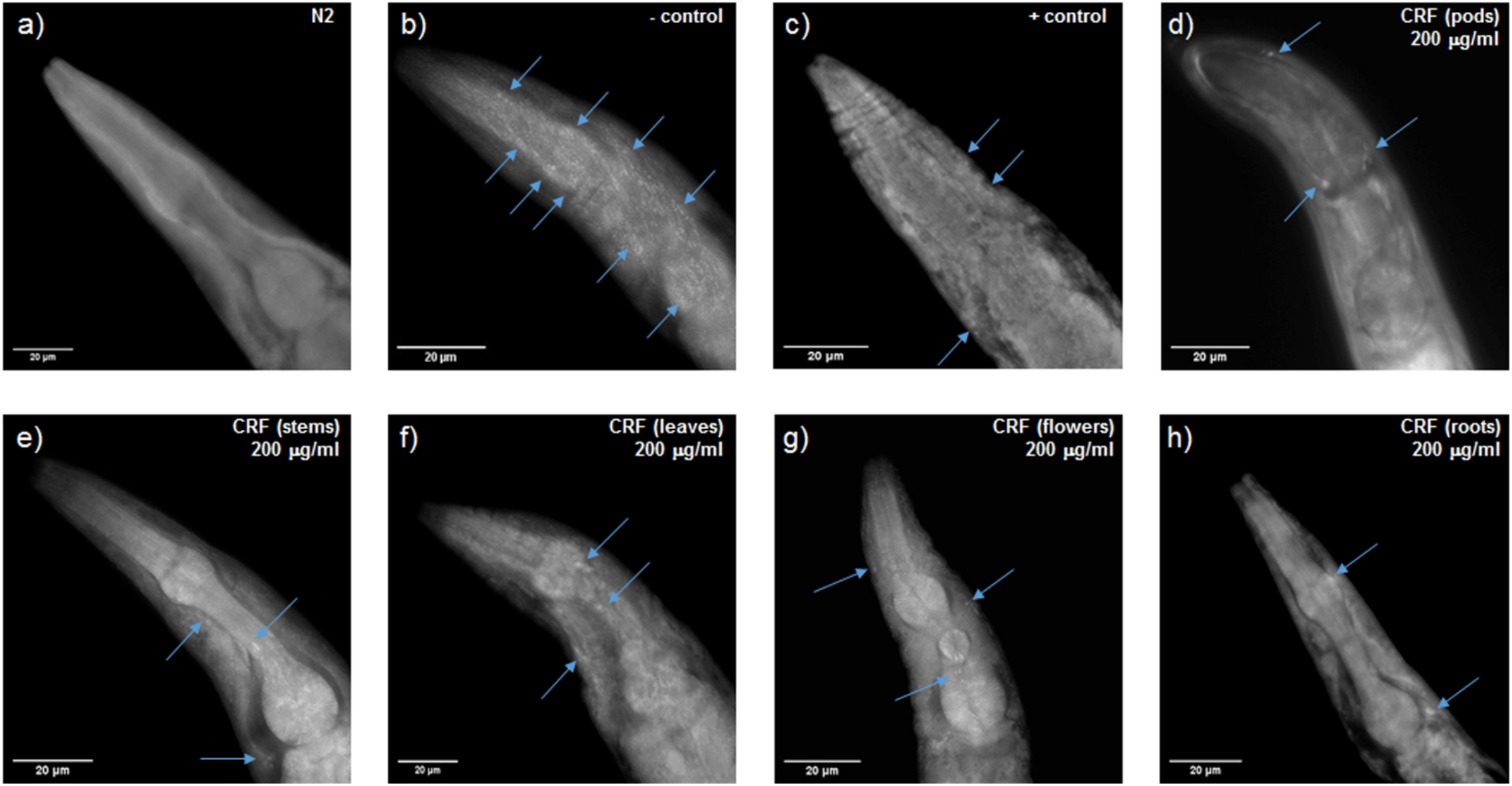
Representative images of Aβ plaques in the head region of transgenic strain CL2006. Age-synchronized L1 worms were spotted on plates with OP50 at 20 °C for 72 h. Worms were then transferred on NGM plates and treated with vehicle (negative control) or CRFs (concentration: 200 μg/ml) from different tissues or Vitamin C (positive control) for 48 h. Worms were then fixed, permeabilized and stained with 0.125% thioflavin-S. Representative images of (a) wild type (b) negative control (c) positive control (d) CRF from pods (e) CRF from stems (f) CRF from leaves (g) CRF from flowers and (h) CRF from roots. Arrows indicate Aβ deposits. Scale bars: 20 μm.

### Molecular docking of cyclotide and Aβ structures

In order to obtain a structural understanding of how the presence of cyclotides would reduce the aggregation of Aβ peptides, we carried out three-dimensional modeling of the interactions and molecular dynamics simulations. A general schema for docking and MD simulation of the multi-protein complexes employed in the present study is described in Figure 7. We used five different Aβ peptide structures, including Aβ_1-40_ and Aβ_1-42_, to evaluate the possible interaction modes between Aβ and cyclotides. The Aβ quaternary structures used are: PDB ID 1IYT (Aβ_1-42_ monomer), PDB ID 2BEG, (U-shaped pentamer Aβ_17-42_), PDB ID 2MXU (S-shaped model Aβ_11- 42_), d) PDB ID 2NAO (disease relevant Aβ_1-42_ fibrils) and PDB ID 2M4J (Aβ_1-40_ isolated from brain of AD patient) (Figure 7a) ^31–35^. Only one NMR structure of chemically synthesized Cter-M cyclotide from *C. ternatea* (PDB ID 2LAM) is known till date and we have used it as a representative cyclotide structure for all the docking and simulation studies (Figure 7b) ^36^. We carried out rigid-body docking studies using FRODOCK2.0 between cyclotide and different forms of Aβ peptide ^37^. PPCheck was used to predict the best docking pose based on the total stabilizing energy and normalized energy per residue for each of protein-peptide complex ^38^. For each of the interactions *i*.*e*. 1IYT-2LAM, 2BEG-2LAM, 2MXU-2LAM, 2NAO-2LAM and 2M4J-2LAM complexes, the best ranking pose was selected to perform MD simulations of 100 ns (Table S5, Supporting Information).

**Figure 7.**
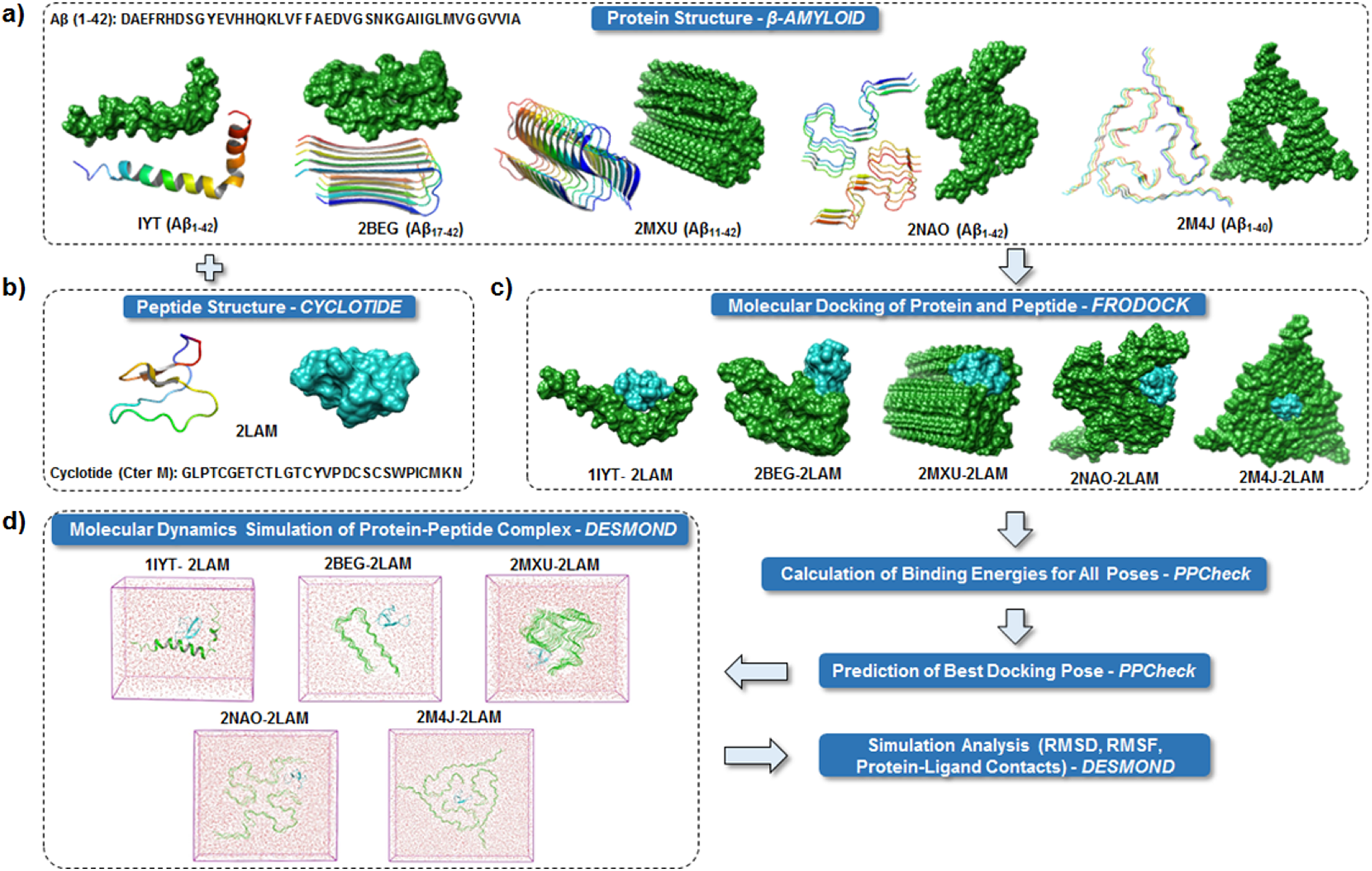
Schematic representation of the protocol used for the docking and MD simulation study. Cartoon and surface representations of the five different Aβ structures. (b) Cartoon and surface representation of cyclotide Cter-M structure. (c) Surface representations of the best binding mode of the five docked complexes of Aβ and cyclotides. Color scheme used for surface representation: green - Aβ and cyan - cyclotide. (d) The protein complexes are minimized and placed in a box of water. The systems are then set up for MD simulation.

### Molecular dynamics simulation analyses of cyclotide - Aβ complexes

To understand the stability of the docked poses for each protein-peptide complex, molecular dynamics simulations were carried out using the Desmond package of Maestro Schrodinger ^39^. The averaged backbone root mean square deviations (RMSD), root mean square fluctuation (RMSF) and radius of gyration (Rg) of the five ensembles of Aβ with and without cyclotides were calculated as a function of time. Between the bound and unbound forms of Aβ, there were greater fluctuations seen in the cyclotide-bound Aβ structure compared to unbound Aβ, based on backbone RMSD and radius of gyration parameters (Figure S2a-e, Supporting Information). As evident from the plot of RMSD variation, all systems experienced some degree of fluctuations at first, but gradually tended to converge after 60 ns implying that the simulations reached equilibrium. PPCheck and Maestro was utilized for identifying all the possible protein-protein interactions including classical hydrogen bonds, aromatic hydrogen bonds, electrostatic and hydrophobic interactions, salt bridges and π-π interactions. The stability of the cyclotide Cter-M (PDB ID: 2LAM) interaction with the disease relevant Aβ_1-42_ fibril (PDB ID: 2NAO) is highlighted in Figure 8. It is evident that upon binding to Cter-M, Aβ_1-42_ fibril undergoes large conformational changes and the secondary structures, especially extended β-sheet conformations, begin to disrupt soon after 25 ns (Supporting Information Movie 1). This is in contrast to the unbound Aβ_1-42_ fibril without the cyclotide molecules wherein extended β-sheet conformations are relatively more stable throughout the simulation (Figure 8 a,b, Supporting Information Movie 1 and 2). The timeline analysis of the secondary structural variations during the 100 ns MD simulations shows that extended β-sheet conformations (in yellow) were stable throughout the trajectory in the unbound Aβ_1-42_ fibril. However, in the cyclotide-Aβ_1-42_ bound trajectory, it is evident that the extended β-sheet conformations undergo structural variations soon after 10 ns, especially around residues 2-7 and 38-41 in each chain (Figure S3, Supporting Information). Figure 8c,d summarizes the molecular interactions between the Aβ_1-42_ fibril and Cter-M at the beginning (0^th^ ns snapshot) and end (100^th^ ns snapshot) of the simulation period, respectively. At t=0 ns, the Aβ_1-42_ fibril and Cter-M complex is mainly stabilized by nine hydrogen bonds, five hydrophobic interactions and two electrostatic interactions (Table 3; Figure 8c). The interaction interface involves polar and aromatic residues of Aβ *i*.*e*., His13, His14 from chain B, Phe4, His6, Tyr10, His13 and His14 from chain C and aliphatic residues such as Val40 from chain D, Gly38 and Val40 from chain E, and Gly38, Val39 and Val40 from chain F. The residues of Cter-M involved in the interaction are Gly1, Pro3, Gly6, Gly7, Tyr15, Val16, Ile25, Cys26, and Asn29. At t=100 ns, the Aβ_1-42_ fibril and Cter-M complex is mainly stabilized by six hydrogen bonds, six hydrophobic interactions and three electrostatic interactions (Table 3; Figure 8d). The interaction interface at the end of 100ns simulations involves similar residues of Aβ as mentioned earlier *i*.*e*., His14 from chain B, Glu3, Phe4, His6, Tyr10, His13 and His14 from chain C, and aliphatic residue such as Val40 from chain D, E and F. Cter-M residues stabilizing the Aβ_1-42_ are also comparable comprising of Gly1, Leu2, Pro3, Gly6, Gly7, Thr8, Tyr15, Val16, Lys28 and Asn29. Figure S4 shows that cyclotide has the potential to stably bind and disrupt the ordered structures of several forms of amyloid fibrils and the interactions remain stable throughout (Figure S4, Supporting Information). The intermolecular hydrogen bonds in all the five protein-peptide complexes were also monitored throughout the simulation period as these are relative measures of binding affinity. The average number of hydrogen bonds between complexes 1IYT-2LAM, 2BEG-2LAM, 2MXU-2LAM, 2NAO-2LAM and 2M4J-2LAM were 3, 1, 3, 4 and 3, respectively (Figure S2, Supporting Information). The total stabilizing energy at t=0 ns and t=100 ns frames for each of the cyclotide - Aβ complexes is detailed in Table S6 (Supporting Information). Interactions between the other four pairs of Aβ and cyclotides are illustrated in Figure S5 and Tables S7-S10 (Supporting Information).

**Table 3.** Analysis of protein-protein interactions at t=0 ns (top) and t=100ns (bottom) frames between cyclotide Cter-M (PDB ID: 2LAM) and Aβ_1-42_ fibril (PDB ID: 2NAO).

| Interactions | <sup>a</sup>Maestro | <sup>a</sup>PPCheck | 2NAO |  |  |  | 2LAM |  |  |  | <sup>b</sup>Type<br>of<br>Bond | Distance<br>Å |
| --- | --- | --- | --- | --- | --- | --- | --- | --- | --- | --- | --- | --- |
|  |  |  | Residue-1 |  |  |  | Residue-2 |  |  |  |  |  |
|  |  |  | Res<br>No. | Res<br>Name | Chain | Atom<br>Name | Res<br>No. | Res<br>Name | Chain | Atom<br>Name |  |  |

| t=0 ns |  |  |  |  |  |  |  |  |  |  |  |  |
| --- | --- | --- | --- | --- | --- | --- | --- | --- | --- | --- | --- | --- |
| Hydrogen Bond | ✓ | - | 13 | His | C | NE2 | 7 | Glu | Z | N | SB | 3.2 |
|  | ✓ | - | 13 | His | B | ND1 | 3 | Pro | Z | O | SB | 3.7 |
|  | ✓ | - | 14 | His | B | NE2 | 6 | Glu | Z | N | SB | 3.5 |
|  | ✓ | ✓ | 38 | Gly | E | O | 15 | Tyr | Z | OH | BS | 2.7 |
|  | - | ✓ | 39 | Val | F | N | 15 | Tyr | Z | OH | BS | 2.8 |
|  | - | ✓ | 38 | Gly | F | N | 15 | Tyr | Z | OH | BS | 2.9 |
| Aromatic Hydrogen Bond | ✓ | - | 4 | Phe | C | CZ | 29 | Asn | Z | O | SB | 3.8 |
|  | ✓ | - | 6 | His | C | CE1 | 1 | Gly | Z | O | SB | 4.4 |
|  | ✓ | - | 13 | His | C | CE1 | 26 | Cys | Z | O | SB | 3.2 |
| Hydrophobic Interactions | - | ✓ | 10 | Tyr | C | CB | 25 | Ile | Z | CB | SS | 5.0 |
|  | - | ✓ | 40 | Val | D | CB | 16 | Val | Z | CB | SS | 6.9 |
|  | - | ✓ | 40 | Val | E | CB | 15 | Tyr | Z | CB | SS | 6.5 |
|  | - | ✓ | 40 | Val | E | CB | 16 | Val | Z | CB | SS | 6.6 |
|  | - | ✓ | 40 | Val | F | CB | 15 | Tyr | Z | CB | SS | 5.3 |
| Electrostatic Interactions | - | ✓ | 13 | His | C | CB | 7 | Glu | Z | CB | SS | 7.9 |
|  | - | ✓ | 14 | His | C | CB | 7 | Glu | Z | CB | SS | 7.6 |
| t=100 ns |  |  |  |  |  |  |  |  |  |  |  |  |
| Hydrogen Bond | ✓ | ✓ | 6 | His | C | N | 1 | Gly | Z | O | BB | 2.9 |
|  | ✓ | ✓ | 4 | Phe | C | N | 29 | Asn | Z | O | BB | 2.8 |
|  | ✓ | - | 14 | His | C | ND1 | 8 | Thr | Z | OG1 | SS | 2.8 |
| Aromatic Hydrogen Bond | ✓ | - | 6 | His | C | CD2 | 1 | Gly | Z | O | SB | 3.2 |
|  | ✓ | - | 14 | His | B | CE1 | 6 | Gly | Z | O | SB | 3.5 |
| Hydrophobic Interactions | - | ✓ | 4 | Phe | C | CB | 2 | Leu | Z | CB | SS | 4.6 |
|  | - | ✓ | 10 | Tyr | C | CB | 25 | Ile | Z | CB | SS | 6.8 |
|  | - | ✓ | 40 | Val | D | CB | 15 | Tyr | Z | CB | SS | 5.5 |
|  | - | ✓ | 40 | Val | D | CB | 16 | Val | Z | CB | SS | 5.7 |
|  | - | ✓ | 40 | Val | E | CB | 15 | Tyr | Z | CB | SS | 5.1 |
|  | - | ✓ | 40 | Val | F | CB | 15 | Tyr | Z | CB | SS | 7.0 |
| Electrostatic Interactions | - | ✓ | 3 | Glu | C | CB | 28 | Lys | Z | CB | SS | 9.4 |
|  | - | ✓ | 13 | His | C | CB | 7 | Glu | Z | CB | SS | 9.1 |
|  | - | ✓ | 14 | His | C | CB | 7 | Glu | Z | CB | SS | 7.5 |

<sup>a</sup> Protein-protein interaction identified from Maestro and/or PPCheck is highlighted with ✓.
<sup>b</sup> "SS" represents sidechain-sidechain interaction, "SB" represents sidechain-backbone interaction, and "BB" represents backbone-backbone mode of interaction between the two interacting amino acids.

**Figure 8.**
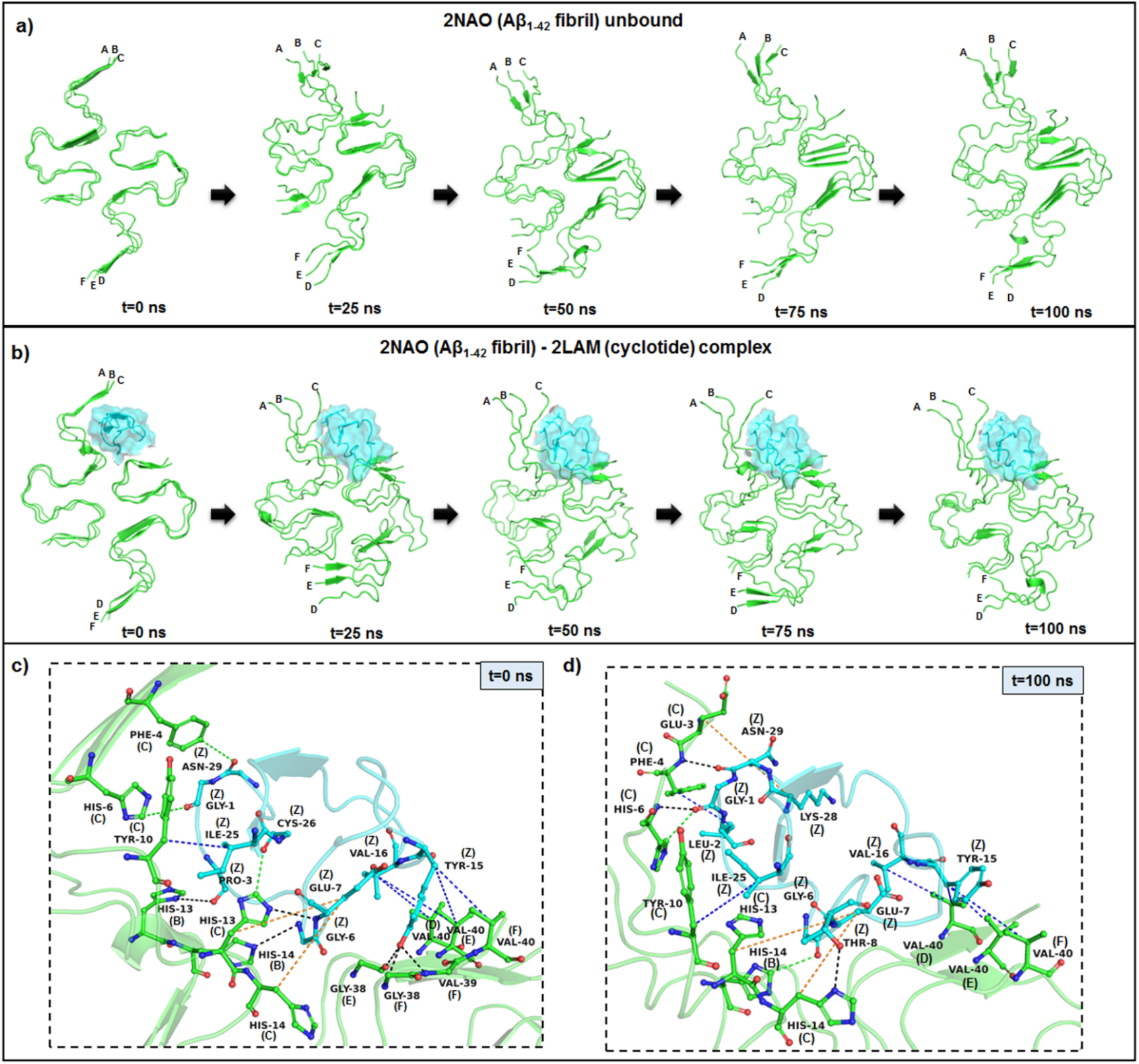
Representative example of MD simulations between the Aβ_1-42_ fibril and cyclotide Cter-M. (a) Snapshots of MD simulation of unbound Aβ_1-42_ fibril (PDB ID: 2NAO; green colored cartoon representation) at different time points along the simulation period. (b) Snapshots of MD simulation of Aβ_1-42_ fibril (PDB ID: 2NAO; green colored cartoon representation) and Cter-M (PDB ID: 2LAM; cyan surface representation) complex at different time points along the simulation period. The cyclizing residues (Gly 1 and Asn 29) are shown as sticks to indicate the N- and C-termini. Capital letters, which denote the polypeptide chains of 2NAO, are placed at the 1^st^ residue of each chain. (c,d) Molecular interactions between the Aβ_1-42_ fibril (PDB ID: 2NAO; chains A-F; green cartoon representation) and cyclotide Cter-M (PDB ID: 2LAM; chain Z; cyan cartoon representation) at the beginning (0th ns snapshot) and end (100th ns snapshot) of the simulation period, respectively. Color scheme for interactions used: classical hydrogen bonds in black, aromatic hydrogen bonds in green, hydrophobic interactions in blue, electrostatic interactions in orange, salt-bridges in pink and π-π interactions in red.

## DISCUSSION & CONCLUSIONS

This study investigates the potential of cyclotides from *C. ternatea* as an inhibitor of Aβ aggregation using *C. elegans*. Synthetically created cyclotides, analogous to the *Oldenlandia affinis* plant-derived peptide, have been shown to suppress multiple sclerosis (MS) when administered orally ^40^. More recently, anti-neurodegenerative properties of cylotides from *Psychotria solitudinum* were reported to act as inhibitors of human prolyl oligopeptidase, a promising target for the treatment of cognitive deficits in several psychiatric and neurodegenerative diseases ^41^. However, up till now there have been no reports of pharmacological activities of cyclotides derived from *C. ternatea* for the treatment of neurodegenerative disorders such as AD, Parkinson’s disease or Huntington’s disease. In the present study, we isolated CRF from five tissues of *C. ternatea* and identified 67 cyclotide-like masses from MS experiments, of which 22 were sequenced using the available nucleotide-level information. Using CL4176 strain, we tested the effects of different CRF concentrations from various tissues. Our data show a significant delay in Aβ induced paralysis in nematodes treated with CRF, irrespective of the plant tissue, and the effects were concentration-dependent underscoring the neuroprotective effects of cyclotides. The lower doses of CRF tested (20 and 80 μg/ml) produced less significant protective effects, however, the highest dose tested (200 μg/ml) delayed the paralysis phenotype drastically compared to the untreated and positive controls. Many compounds such as scorpion venom peptides ^22^, sesamin and sesamolin ^42^, and tryptophan-containing heptapeptides ^24^ exert protective effects against Aβ toxicity tested using the same strain of *C. elegans*. With the development of newer transgenic *C. elegans* models, Aβ expression has been made to express pan-neuronally ^43^. CL2355 strain has inducible neuronal Aβ_1-42_ expression which leads to defects in 5-HT sensitivity, chemotaxis behavior and altered movement in liquid ^43^. Here, under normal conditions, CL2355 nematodes exhibit chemotaxis dysfunction to volatile cues, but upon CRF treatment, chemotaxis was rescued to a reasonable extent indicating protection against Aβ induced toxicity in the neuronal cells. The lowest dose of CRF tested (20 μg/ml) was not sufficient to alleviate Aβ toxicity, however, higher doses (80 and 200 μg/ml) improved the rescue of chemotaxis defects. The observed effect could also be due to the presence of other molecules in the CRF, however, we speculate that the effect might be mainly due to cyclotides as CRF contains predominantly cyclotides. Teasing apart the role of individual cyclotide from CRF warrants further investigation given the multiple challenges with separation and characterization ^44^.

Oxidative stress is theorized to be a major contributing factor to AD as it is induced by β-amyloid aggregation ^45-47^. Several evidences correlate AD pathogenesis with increased oxidative stress resulting from increased ROS. Earlier studies have reported that cyclotides exhibit antioxidant properties ^48-50^, and our findings are in line with these reports. Interestingly, feeding CL4176 worms with CRF from flowers of *C. ternatea* exhibited the most significant reduction of intracellular ROS. It has been reported earlier that in cysteine-stabilized peptides, the cysteine and methionine residues have antioxidant activities as they can directly react with free radicals ^49,51^. Here, the six cysteine residues in cyclotides could be contributing to the antioxidant property. Next, using CL2006, we qualitatively assessed the amount of Aβ deposition in the neurons of worms treated with CRF and observed that the deposits were reduced after treatment with the highest dose in line with earlier observations.

Our results clearly indicate that the structural scaffold of cyclotide is more important than sequences as CRF activity is tissue-specific. Aβ is an extracellular protein and in its monomeric form, in the membrane adopts an α-helical structure and conformationally transitions into β-sheet rich structures in the process of aggregation ^52^. Here, we show that the cyclotide interaction with Aβ fibrils can be reinforced by hydrogen bonding, hydrophobic and long-range electrostatic interactions between key residues of Aβ peptide and cyclotide. We show a representative complex of disease-relevant Aβ_1-42_ fibril (PDB ID: 2NAO) and cyclotide Cter-M (PDB ID: 2LAM) to highlight the strong interactions between the two molecules. It is clearly evident that the protein-peptide pair display several non-covalent interactions throughout the simulation period and the nature of residue interactions are highly conserved. The persistence of high number of intermolecular hydrogen bonds and strong non-covalent interactions throughout the trajectory highlights the stable binding of cyclotide in the exposed pockets of amyloid fibril. Moreover, cyclotide disrupts the inter-chain hydrogen bonds and salt bridges in Aβ which are crucial for the fibril structure and shape.

In conclusion, our study reports 22 cyclotide sequences in the CRFs using a combined transcriptomics and proteomics approach and provides a systematic activity evaluation of cyclotides identified. We show that CRFs can (1) attenuate Aβ-induced paralysis in the muscle cells of transgenic CL4176 worms, (2) rescue chemosensory defects associated with Aβ-expression in the neuronal cells of transgenic CL2355 worms, (3) diminish ROS production caused by oxidative stress in transgenic CL4176 worms and (4) reduce Aβ-plaques and oligomers in transgenic CL2006 worms. Further, molecular docking and MD simulations model the mode of interaction between cyclotides and different physiological forms of Aβ structures. Amongst the 22 cyclotides identified in this study, we performed computational analyses only using Cter M, the most abundant cyclotide from *C. ternatea* whose structure is available. Using this as a representative, we have described possible binding events with Aβ aggregates. Taken together, our findings highlight the neuroprotective effects of cyclotides from *C. ternatea* and their immense potential in peptide therapeutics as Aβ inhibitors.

## EXPERIMENTAL SECTION

### Extraction of cyclotides from *C. ternatea*

Plant collection and cyclotide extraction were performed using established methods. Briefly, fresh plant parts such as pods, stems, leaves, flowers and roots were collected and oven dried at 70 °C. Each tissue was extracted with 1:1 DCM: MeOH (v/v) with overnight stirring. The extract was partitioned with 1:1 mixture of DCM: (MeOH: ddH_2_O, 2:3) and fractionated on a packed RP-C_18_ silica gel column (0%–100% gradient elution using ACN/ddH_2_O) (particle size: 40-63 μm; pore size: 60 Å; Polygosil®). Each eluate was examined using MALDI-TOF (UltraFlex Bruker Daltonics) for cyclotide-like masses (2.5-4.0 kDa). Fractions eluted with 50, 70 and 100% ACN/ddH_2_O showed cyclotide-like masses). These fractions were pooled for each plant tissue and freeze-dried (referred henceforth as “crude extracts”) for further RP-HPLC purification.

### Purification and MALDI-TOF analysis

HPLC (Shimadzu Prominence) purification of crude extracts was performed on a semi-preparative Phenomenex Proteo C_18_ column (250 × 10 mm, 10 μm, 110 Å) at flow-rate of 3 ml/min with a fraction collector. A linear gradient of 1% min^-1^ of 0–95% buffer B (100% acetonitrile, 0.1% trifluoroacetic acid) was applied, and the eluents were monitored at 220, 254, and 280 nm. Early and late-eluting peaks were separated into five fractions (A-E), manually collected for each plant tissue and lyophilized. Each of the five fractions, per tissue, was subjected to MALDI-TOF mass-spectrometry (UltraFlex Bruker Daltonics) analysis with 50 Hz pulsed nitrogen laser (λ=337 nm), in positive ion reflectron mode. The samples were prepared in 1:1 mixture of H_2_O-ACN with 0.1% TFA by mixing an equal amount of peptide fractions (0.5 μl) with α-cyano-4-hydroxycinnamic acid (CCA)/2,5-dihydroxy benzoic acid (DHB) matrix (Sigma) and spotted on a stainless-steel plate and air-dried. HPLC fraction D from the crude extracts of each of the five tissues showed maximum cyclotide-like masses (2.5-4.0 kDa), both in terms of number and abundance, and fewer small molecule interferences. Total protein content in the lyophilized fraction D (termed as cyclotide-rich fraction (CRF) henceforth) - was determined by Bradford protein assay based on standard protocol. Stock solutions of CRF were prepared by dissolving the lyophilized CRF powder in 70% ACN (∼5 mg/ml) and stored at −20 °C until use. For the assays, dilutions were prepared from the stock solutions to obtain three concentrations of CRF *i*.*e*. 20, 80, 200 μg/ml.

### Combined -omics approach for cyclotide identification

Cyclotide precursor and mature sequences were identified using the previously reported transcriptome assembly ^25^ and their monoisotopic masses were calculated (Table S1, Supporting Information). These calculated monoisotopic masses were then compared to the observed monoisotopic masses from MALDI-TOF spectra of CRFs from five tissues (pods, stems, leaves, flowers and roots).

### Worm strains and maintenance

*C. elegans* strains N2, CL4176, CL2006 and CL2355 were a kind gift from Dr. Ashwini Godbole (The University of Trans-Disciplinary Health Sciences and Technology). The transgenic strains expressing human Aβ used in the current study are detailed in Table 1. Wild type *C. elegans* strain N2 (Bristol) were grown and maintained on standard nematode growth medium (NGM) seeded with *Escherichia coli* strain OP50 as a food source and maintained at 20 °C. The transgenic CL4176 (smg-1 (cc546); myo-3::Aβ_1–42_::3′-UTR (long)), CL2006 (unc-54::Aβ_1–42_ (wild type); dimer Aβ_1–42_ or Met^35^Cys Aβ_1–42_) and CL2355 (smg-1 (cc546); snb-1::Aβ_1-42_::3’-UTR (long) + mtl-2::GFP) strains were grown and maintained on standard NGM media seeded with *E. coli* strain OP50 and maintained at 15-16 °C. To obtain age-synchronized worms, the reproductive worms (3 days of age) were transferred to the fresh NGM plates and allowed to lay eggs for 4-6 h. Worms hatched from these eggs were used for the study.

### Paralysis assay

Paralysis assays were performed as shown in Figure 3a. Briefly, age-synchronized transgenic CL4176 strains were maintained at 16 °C on NGM plates (60 x 10 mm culture plates; ∼20 eggs/plate) spotted with OP50 containing vehicle, *i*.*e*. 100 mM ammonium bicarbonate buffer (negative control) and treated with or without CRF from different tissues (concentrations: 20, 80, 200 μg/ml). Vitamin C (0.1 μg/ml) was used as a positive control. Transgenic expression of Aβ in body wall muscles in CL4176 strains can be induced by temperature upshift from 16 °C to 23 °C (starting at 36 h after egg laying). Paralysis was scored every 8 hours, starting from the 20th hour until ∼80 % of the worms in the negative control experiment were paralyzed. The worm was considered paralyzed if it did not move or only head movement was observed upon prodding with a platinum loop. Paralysis time course was plotted.

### Chemotaxis assay

Chemotaxis assays were performed as described in Figure 4a. Briefly, synchronized transgenic CL2355 strains were maintained at 16 °C on NGM plates (90 x 15 mm culture plates). They were spotted with OP50 containing vehicle (100 mM ammonium bicarbonate buffer) and with or without CRF from different tissues (concentrations: 20, 80, 200 μg/ml). Vitamin C (0.1 μg/ml) was used as a positive control. The assay was started from eggs maintained at 16 °C for 36 h (∼20-60 eggs/plate), and then temperature was upshifted to 23-25 °C to induce expression of neuronal Aβ and incubated for another 36 h. Worms were then collected, washed with M9 buffer thrice, assayed on 90 mm plates containing 1.9% agar, 1 mM CaCl_2_, 1 mM MgSO_4_ and 25 mM phosphate saline buffer. Worms were placed at the center, 2 μl of 0.1% benzaldehyde in 100% ethanol and 1 μl sodium azide were added at positions A & D as attractant odorants (Figure 4a). 2 μl of 100% ethanol and 1 μl sodium azide were added at positions B & C as control odorants. Assay plates were incubated at 23-25 °C for 2 hours and Chemotaxis Index (C.I.) was calculated using the formula: C.I. = [(Number of worms at attractant position - numbers of worms at control position)/Total number of worms scored)].

### Intracellular ROS assay

ROS levels were analyzed in whole live nematodes using cell-permeable molecular probe, H_2_DCF–DA. Inside the cell, deacetylated by the intracellular esterases, H_2_DCF–DA is retained whereas upon oxidation by ROS, non-fluorescent H_2_DCFDA is converted into highly fluorescent 2’,7’-dichlorofluorescein (DCF). The intensity of DCF fluorescence correlates with the levels of intracellular ROS. For detection of ROS production, age-synchronized *C. elegans* (CL4176; 100 eggs per plate) were treated with or without CRF from different tissues (concentrations: 200 μg/ml) for 36 h at 16°C followed by a shift to 23 °C for 36 h. The worms were collected and the *E. coli* (OP50) bacteria was removed by washing thrice with PBS-T. A 200 μl volume of the suspension was pipetted into a 96-well plate. Fresh 100 μM H_2_DCF–DA solution was prepared in PBS-T from 50 mM stock solution. 50 μl of 100 μM H_2_DCF–DA was then added to each well. Immediately after the addition, the basal fluorescence of each well was measured in a microplate reader at excitation/emission wavelengths of 485 and 538 nm. The plates were read at 37 °C every 30 min for 2.5 h. The time-dependent increase in the fluorescence was linear in this time period. We therefore chose to use fluorescence intensity at 180 min for the quantification. Vitamin C (0.1 μg/ml) was used as a positive control. To further examine the changes in ROS the worms from the plate were observed at 10x magnification using a fluorescence microscope (DM5000B, Leica, Germany) with an FITC excitation and emission filter setup. The images were acquired using a CCD digital camera (Hamamatsu ORCA-ER).

### Fluorescence staining of β-amyloid

Age-synchronized CL2006 were plated on NGM plates (60 x 10 mm culture plates; ∼30 eggs/plate) spotted with OP50 at 20 °C for 72 h. Then worms were transferred on NGM plates containing vehicle and treated with or without CRF from different tissues (concentrations: 200 μg/ml), for 48 h. Vitamin C (0.1 μg/ml) was used as a positive control. The worms were then fixed in 4% paraformaldehyde in phosphate-buffered saline (PBS), at pH 7.4 in 4 °C for 24 h. The worms were then permeabilized in 1% Triton X-100, 5% fresh β-mercaptoethanol and 125 mM Tris, at pH 7.4, and incubated at 37 °C for another 24 h. The worms were washed 2–3 times with PBS-T and then stained with 0.125% thioflavin-S (Sigma) in 50% ethanol for 2 min and then de-stained sequentially in ethanol (50%, 75% and 90%). The worms were mounted on coverslips covered with 80% glycerol and observed under a fluorescent microscope (DM5000B, Leica, Germany) for the presence of amyloid plaques in the head region of individual worms. The images were acquired at 40x magnification using a CCD digital camera (Hamamatsu ORCA-ER).

### Protein - Peptide docking

In order to better understand and visualize the interactions between different forms of Aβ fibrillar species and cyclotides, we performed molecular docking experiments using FRODOCK2.0 ^37^. The five three-dimensional NMR solution structures (first model amongst the ensembles) of Aβ used in this study are: a) PDB ID: 1IYT, Aβ_1-42_ monomer, b) PDB ID: 2BEG, U-shaped pentamer Aβ_17-42_, c) PDB ID: 2MXU, S-shaped model Aβ_11-42_, d) PDB ID: 2NAO, disease relevant Aβ_1-42_ fibrils and e) PDB ID: 2M4J, Aβ_1-40_ isolated from brain of AD patient. We used the three-dimensional NMR solution structure (first model amongst the ensembles) of Cter-M cyclotide from *C. ternatea* (PDB ID: 2LAM) to dock against each of the five Aβ fibrillar species. For all the generated docked poses for each of the cyclotide-Aβ complexes (5 complexes), PPCheck ^38^ was used to calculate binding energies and to predict best native-like docking pose (Table S5, Supporting Information).

### Molecular Dynamics Simulation

To confirm stable binding of each Aβ-cyclotide complex, MD simulation without constraints was performed using the Desmond package of Maestro Schrodinger ^39^. The TIP4P solvent model was used with an orthorhombic box shape and buffer distance of 10 Å. The OPLS3e force field was used for building of the system. The system was minimized using steepest descent for 2000 steps until a gradient threshold of 50 kcal mol^−1^ Å^−1^ was reached. The system was neutralized by adding Na^+^ ions and 0.15 M salt (NaCl). The output from the system builder was used for MD simulations. The simulation time was set at 100 ns in the NPT ensemble class. The RESPA integrator was used with a time step of 2.0 fs ^53^. The temperature and pressure were set at 300K and 1 bar, using the Nose-Hoover chain ^54^ and the Martyna-Tobias-Klein method ^55^ respectively. For stability analyses, the entire range of simulation time was considered. RMSD, RMSF, radius of gyration and number of hydrogen bonds was calculated for MD runs of all the five Aβ (protein) and cyclotide (peptide) complexes. A set of control MD runs were performed on each of the unbound Aβ structures. Hydrogen bonds were identified using the Simulation Event Analysis module implemented in the Desmond package. Protein RMSF shows the fluctuation of Aβ residues and peptide RMSF shows fluctuation of cyclotide residues. Protein-peptide interactions were also monitored throughout the simulation time and visualized using PyMOL (The PyMOL Molecular Graphics System, version 2.4.1 Schrödinger, LLC).

### Statistical analysis

Statistical analyses were carried out using GraphPad Prism 7 software package (GraphPad Software, La Jolla, CA, USA). Data was tested for normality and homoscedasticity, whenever required, using D’Agostino-Pearson normality test and Bartlett’s test, respectively. Paralysis, Chemotaxis and ROS assay data were subjected to one-way ANOVA followed by Tukey’s honestly significant difference (HSD) procedure when comparing untreated and CRF treated groups. Differences between positive control and CRF treated groups were analyzed for statistical significance by unpaired two-tailed Student’s *t* test. Further details of statistical analyses of all the assays are given in the Supporting Information.

## ASSOCIATED CONTENT

### Supporting Information

**Figure S1:** MALDI-TOF spectra of HPLC purified fractions A, B, C and E from crude extracts of different tissues (pods, stems, leaves, flowers and roots) of *C. ternatea*.

**Figure S2:** Plots of RMSD (top left panel), RMSF (top right panel), Radius of gyration (bottom left panel) and number of hydrogen bonds (bottom right panel) for complexes between cyclotide (PDB ID: 2LAM) and (a) Aβ_1-42_ monomer (PDB ID: 1IYT), (b) Aβ_17-42_ U-shaped pentamer (PDB ID: 2BEG), (c) Aβ_11-42_ S-shaped model (PDB ID: 2MXU), (d) Aβ_1-42_ disease relevant Aβ_1-42_ fibrils (PDB ID: 2NAO) (e) Aβ_1-40_ isolated from brain of AD patient (PDB ID: 2M4J), along the simulation period.

**Figure S3:** Secondary structure timeline analysis during 100 ns MD simulation of (a) Aβ_1-42_ fibril (PDB ID: 2NAO) bound to cyclotide Cter-M (b) unbound Aβ_1-42_ fibril (PDB ID: 2NAO). Colour code explanation of the secondary structures are shown in the small panel below. T denotes turn (aqua); E represents the β-sheet (yellow); B represents isolated bridges (dark yellow); H denotes α-helix (pink); G indicates the 3_10_ helix (blue); I denotes π-helix (red) and C indicates random coils (white). These structural analyses were computed using VMD timeline plugin.

**Figure S4:** Snapshots of MD simulation of cyclotide (PDB ID: 2LAM; cyan surface representation) and (a) Aβ_1-42_ monomer (PDB ID: 1IYT, green cartoon representation), (b) Aβ_17-42_ U-shaped pentamer (PDB ID: 2BEG, green cartoon representation), (c) Aβ_11-42_ S-shaped model (PDB ID: 2MXU, green cartoon representation), (d) Aβ_1-40_ isolated from brain of AD patient (PDB ID: 2M4J, green cartoon representation) complexes at different time points along the simulation period.

**Figure S5:** Molecular interactions between cyclotide (PDB ID: 2LAM; chain Z; cyan cartoon representation) and (a) Aβ_1-42_ monomer (PDB ID: 1IYT; chain A; green (b) Aβ_17-42_ U-shaped pentamer (PDB ID: 2BEG; chain A-E; green cartoon representation), (c) Aβ_11-42_ S-shaped model (PDB ID: 2MXU; chain A-L; green cartoon representation), (d) Aβ_1-40_ isolated from brain of AD patient (PDB ID: 2M4J; chain A-I; green cartoon representation) at the beginning (0^th^ ns snapshot) and end (100^th^ ns snapshot) of the simulation period. Colour scheme for interactions used: classical hydrogen bonds in black, aromatic hydrogen bonds in green, hydrophobic interactions in blue, electrostatic interactions in orange, salt-bridges in pink and π-π interactions in red.

**Table S1:** *C. ternatea* cyclotide transcript sequences ^25^ and their monoisotopic masses

**Table S2:** Descriptive statistics of paralysis assay in transgenic *C. elegans* CL4176.

**Table S3:** Descriptive statistics of chemotaxis assay in transgenic *C. elegans* CL2355.

**Table S4:** Descriptive statistics of intracellular ROS assay in transgenic *C. elegans* CL2006.

**Table S5:** Results of PPCheck webserver for predicting normalized energy per residue for FRODOCK docked models. Best docking pose for each protein-peptide complex is highlighted in green.

**Table S6:** The total stabilizing energy at t=0 ns and t=100 ns frames for each of the five cyclotide-Aβ complexes.

**Table S7:** Analysis of protein-protein interactions at t=0 ns (top) and t=100ns (bottom) frames between cyclotide (PDB ID: 2LAM) and Aβ_1-42_ monomer (PDB ID: 1IYT).

**Table S8:** PPCheck results for protein-protein interactions at t=0 ns (top) and t=100ns (bottom) frames between cyclotide (PDB ID: 2LAM) and Aβ_17-42_ U-shaped pentamer (PDB ID: 2BEG).

**Table S9:** PPCheck results for protein-protein interactions at t=0 ns (top) and t=100ns (bottom) frames between cyclotide (PDB ID: 2LAM) and Aβ_11-42_ S-shaped model (PDB ID: 2MXU).

**Table S10:** PPCheck results for protein-protein interactions at t=0 ns (top) and t=100ns (bottom) frames between cyclotide (PDB ID: 2LAM) and Aβ_1-40_ isolated from brain of AD patient (PDB ID: 2M4J).

**Movie 1** of MD simulation of Aβ_1-42_ fibril (PDB ID: 2NAO) bound to cyclotide Cter-M (PDB ID: 2LAM).

**Movie 2** of MD simulation of unbound Aβ_1-42_ fibril (PDB ID: 2NAO).

## DECLARATION

All animal experiments performed in the manuscript were conducted in compliance with institutional guidelines.

## ACKNOWLEDGMENT

The authors thank Prof. P. Balaram (National Centre for Biological Sciences, Bangalore, India) for his valuable suggestions throughout the project and help with the initial version of the manuscript. We thank Dr. Ashwini Godbole (The University of Trans-Disciplinary Health Sciences and Technology, Bangalore, India) for *C. elegans* strains and Dr. Varsha Singh (Indian Institute of Science, Bangalore, India) for her inputs on the manuscript. We also thank the *C. elegans* facility at NCBS (TIFR). NK was supported by the Tata Education and Development Trust. Mass spectra was obtained using the Proteomics Facility supported by the Department of Biotechnology at the Indian Institute of Science, Bangalore. We thank NCBS (TIFR) and JC Bose fellowship of RS (SB/S2/JC-071/2015). Funding to RV by DST-SERB (Early Career Award, Ramanujan Fellowship), Max Planck Society (Partner group program) and Department of Biotechnology (NER Grant) is gratefully acknowledged.

## AUTHOR CONTRIBUTIONS

Conceived and designed experiments: NK and RV. Performed experiments: NK and HH. Data analysis: NK, RS and RV. Manuscript preparation: NK, RV and RS.

## Table of Contents (TOC)/Graphical Abstract

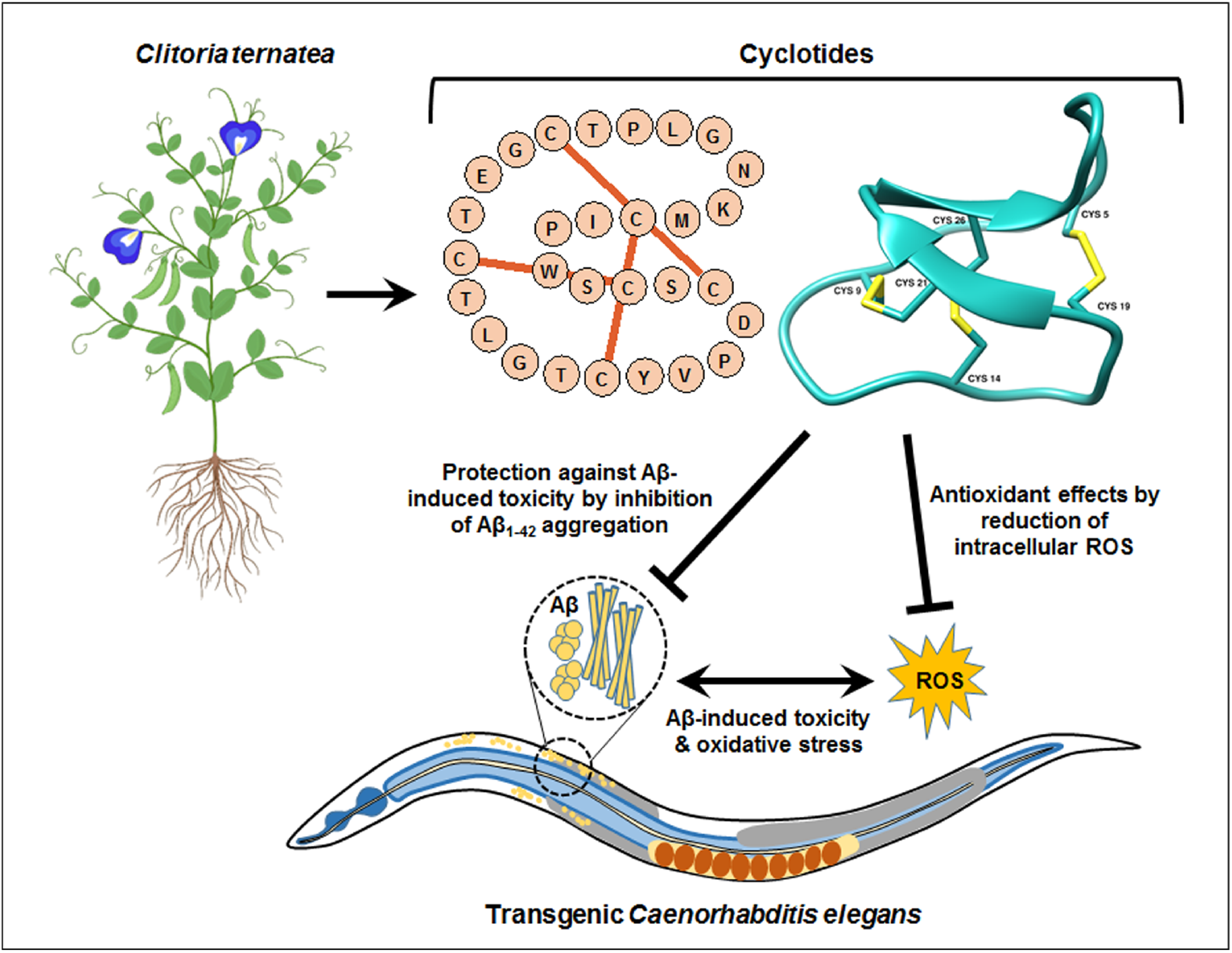

