## Supplemental File for "Disulfide-rich, cyclic peptides from *Clitoria ternatea* protect against β-amyloid toxicity and oxidative stress in transgenic *Caenorhabditis elegans*"

**Figure S1:** MALDI-TOF spectra of HPLC purified fractions A, B, C and E from crude extracts of different tissues a) pods, b) stems, c) leaves, d) flowers and e) roots of *C. ternatea*. The monoisotopic signals (M+= m/z - H+) are labeled.

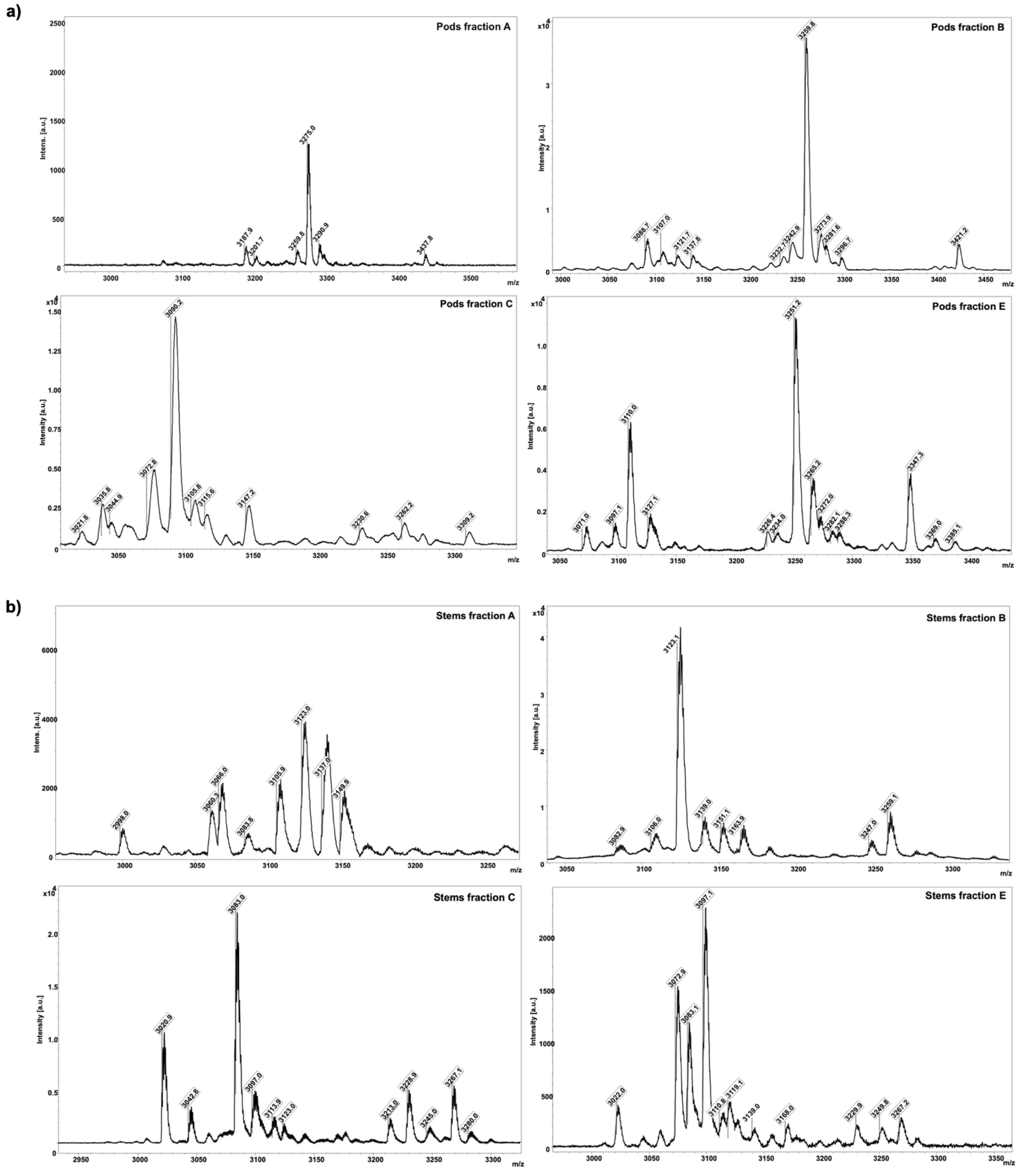

c)

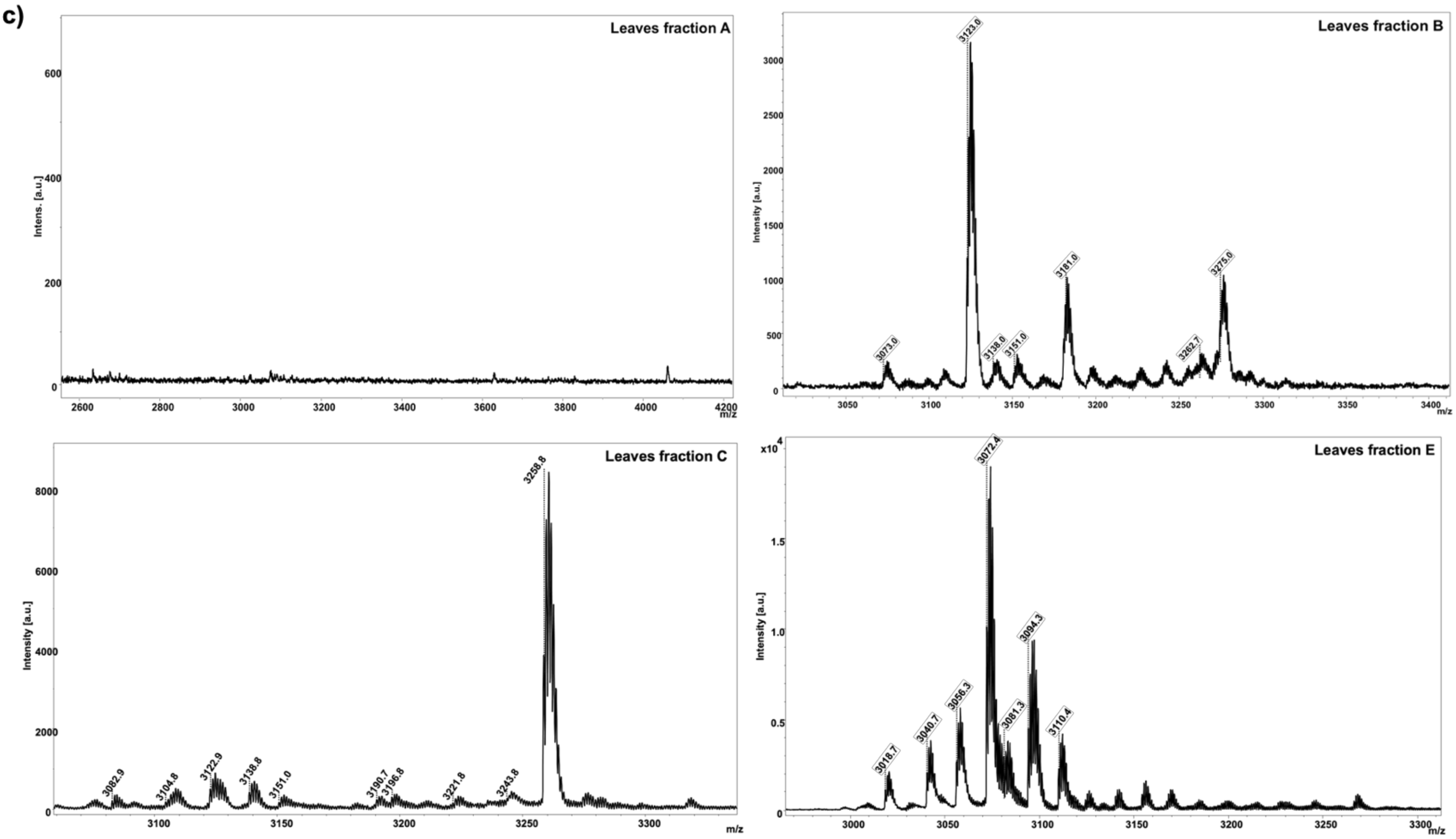
Leaves fraction A
Intens. [a.u.]
m/z
Leaves fraction B
Intensity [a.u.]
3123.0
3073.0
3138.8
3151.0
3181.0
3262.7
3275.0
m/z
Leaves fraction C
Intensity [a.u.]
3258.8
3082.9
3104.8
3122.9
3138.8
3151.0
3190.7
3196.8
3221.8
3243.8
m/z
Leaves fraction E
Intensity [a.u.]
3072.4
3018.7
3040.7
3056.3
3081.3
3094.3
3110.4
m/z

d)

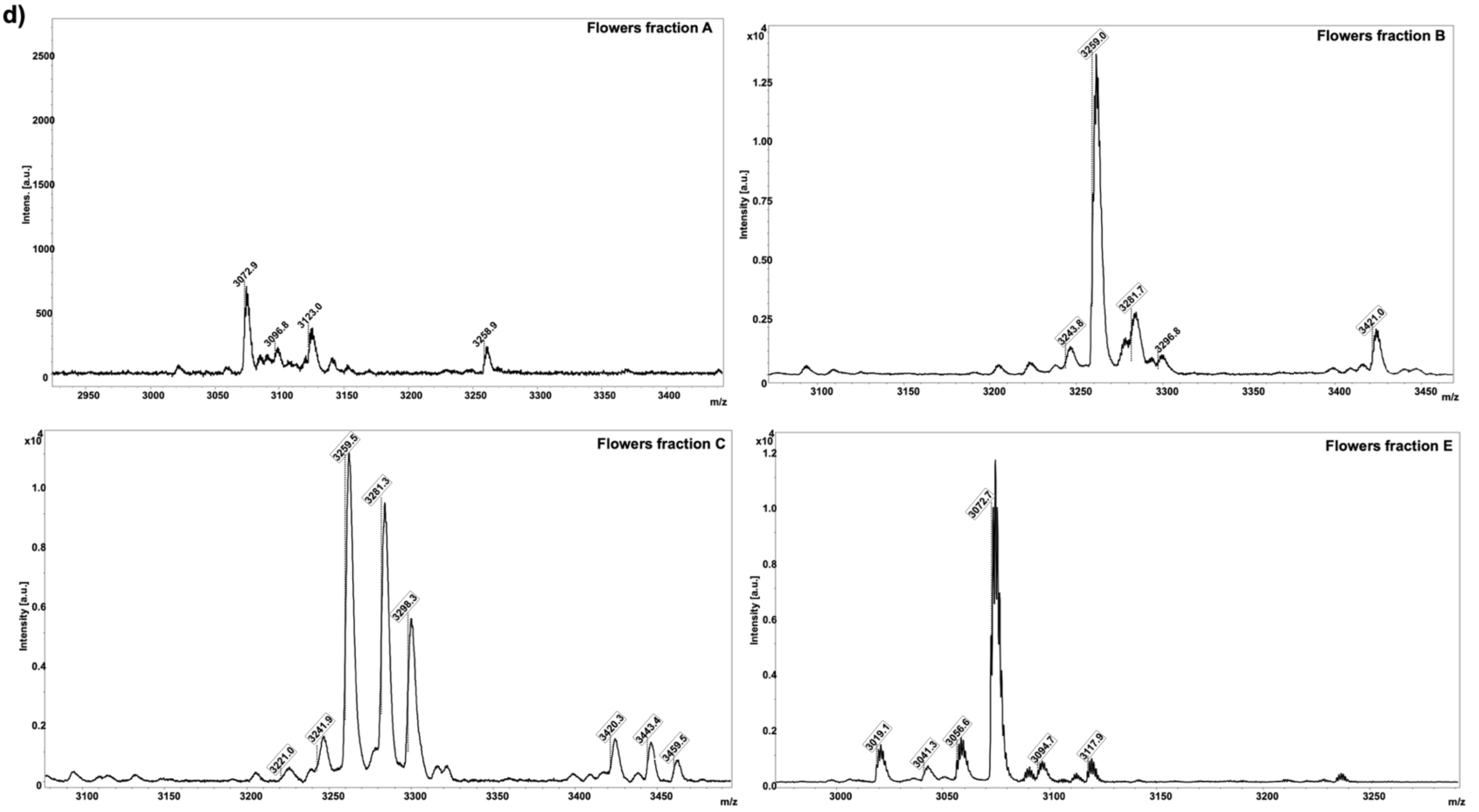
Flowers fraction A
Intens. [a.u.]
3072.9
3096.8
3123.0
3258.9
m/z
Flowers fraction B
Intensity [a.u.]
3259.0
3243.8
3281.7
3296.8
3421.0
m/z
Flowers fraction C
Intensity [a.u.]
3259.5
3281.3
3298.3
3221.0
3241.9
3420.3
3443.4
3459.5
m/z
Flowers fraction E
Intensity [a.u.]
3072.7
3019.1
3041.3
3056.6
3094.7
3117.9
m/z

e)

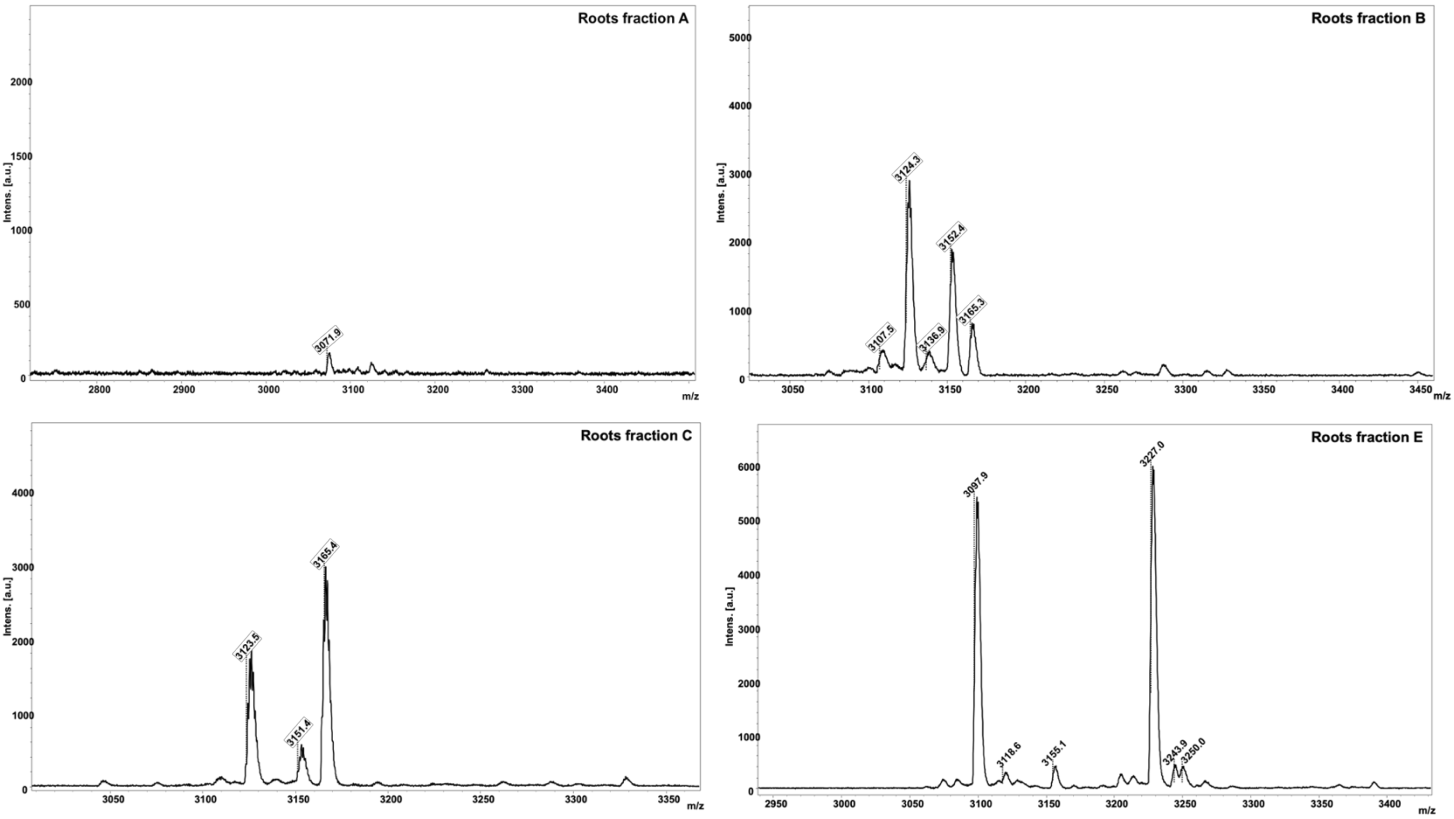

Roots fraction A
Intens. [a.u.]
3071.9
m/z
Roots fraction B
3124.3
3152.4
3107.5
3136.9
3165.3
Roots fraction C
3165.4
3123.5
3151.4
Roots fraction E
3097.9
3227.0
3118.6
3155.1
3243.9
3250.0

**Table S1:** *C. ternatea* cyclotide transcript sequences [25] and their monoisotopic masses

| Sl. No. | GenBank Accession Numbers | Peptide ID | Cyclotide Mature Domain Sequence | Calculated Monoisotopic Mass [Da] | Alternate Names |
|---|---|---|---|---|---|
| 1 | MT468661/ MT468662 | ctr pep 1 | GIPCGESCVFIPCTITALLGCSCKSKVCYKN | 3212.5 | Cter 14; CliotideT40 |
| 2 | MT468663 | ctr pep 2 | GIPCGESCVFIPCLTTVVGCSCKNKVCYNN | 3126.4 | Cliotide T9 |
| 3 | MT468664 | ctr pep 3 | GRPTCGETCFKTKCYTPGCSCSYPICKKN | 3150.4 | |
| 4 | MT468665 | ctr pep 4 | GDALKCGETCFGGTCYTPGCSCDYPICKKN | 3109.3 | |
| 5 | MT468666 | ctr pep 5 | FKTKCYTPGCSCSYPVCKRN | 2262.0 | |
| 6 | MT468667 | ctr pep 6 | GCLPICGETCFKTKCYTKGCSCSYPICKKN | 3253.5 | |
| 7 | MT468668 | ctr pep 7 | GLPICGETCFKTKCYTKGCSCSYPVCKRN | 3164.5 | Cliotide T15; Cter24 |
| 8 | MT468669 | ctr pep 8 | GRPTCGETCFKTKCYTKGCSCSYPVCKRN | 3195.4 | |
| 9 | MT468670 | ctr pep 9 | GNPIVCGETCFFQKCYTPGCSCDAVICTNN | 3162.3 | Cter 25; CliotideT41 |
| 10 | MT468671 | ctr pep 10 | GDPLACGETCFGGTCYTPGCVCDPWPICTKN | 3183.3 | Cter 4; CliotideT39 |
| 11 | MT468672 | ctr pep 11 | ANIPMTCGPCLTDECWTPGCEYHCKYCKNS | 3342.4 | |
| 12 | MT468673/ MT468674 | ctr pep 12 | LPTCGETCTLGTCYVPDCSCSWPICMKN | 3000.2 | |
| 13 | MT468675 | ctr pep 13 | KIPCGESCVWIPCFTSAFGCYCQSKVCYHS | 3321.4 | Cliotide T38 |
| 14 | MT468676/ MT468677/ MT468678 | ctr pep 14 | SCVWIPCITGAIGCSCKNRVCYRN | 2623.2 | |
| 15 | MT468679 | ctr pep 15 | DTTPCGESCVWIPCVSSIVGCSCQNKVCYQN | 3298.4 | Cliotide T13; Cter 23 |
| 16 | MT468680 | ctr pep 16 | GFNSCSEACVYLPCFSKGCSCFKRQCYKN | 3247.4 | Cter 34; CliotideT33 |
| 17 | MT468681 | ctr pep 17 | GVIPCGESCVFIPCISTVIGCSCKNKVCYRN | 3267.6 | Cter A |
| 18 | MT468682 | ctr pep 18 | GISCGESCVFIPCITGIAGCSCKNKVCYLN | 3057.4 | |
| 19 | MT468683 | ctr pep 19 | ESCVWIPCLTGYFGCYCQSKVCYHN | 2880.2 | |
| 20 | MT468684 | ctr pep 20 | CVIPCGESRVFIPCITGAIGCSCKSKVCYRN | 3281.6 | |
| 21 | MT468685 | ctr pep 21 | ARIPCGESCVWIPCTITALVGCACHEK | 2837.3 | |
| 22 | MT468686/ MT468689 | ctr pep 22 | GSITGCGGGCLLGRCYRPGCTCVRRICRRN | 3162.5 | |
| 23 | MT468687/ MT468690 | ctr pep 23 | GSAIRCGERCLLGRCHRPGCTCIRRICRRN | 3390.7 | Cter 13 |
| 24 | MT468688 | ctr pep 24 | GSAIRCGERCLLGRCHRPGCTCVRRICRRN | 3376.7 | |
| 25 | MT468691 | ctr pep 25 | GDLFKCGETCFGGTCYTPGCSCDYPICKNN | 3171.3 | Cliotide T32 |
| 26 | MT468692 | ctr pep 26 | GSSIVTCGETCLRGKCYTPGCTCVRPICKKN | 3252.5 | |
| 27 | MT468693 | ctr pep 27 | GSVIGCGETCLRGRCYTPGCTCDHGICKKN | 3106.4 | Cliotide T16 |

| | | | | | |
|---|---|---|---|---|---|
| 28 | MT468694 | ctr pep 28 | GLPICGETCFTGTCYTPGCTCSYPVCKKN | 3021.3 | Cliotide T18; Cter 6 |
| 29 | MT468695 | ctr pep 29 | GDPFKCGESCFAGKCYTPGCTCSRPICKKN | 3175.4 | |
| 30 | MT468696/ MT468697 | ctr pep 30 | GSPCGESCVFIPCISTVIGCSCKNKVCYRN | 3142.4 | |
| 31 | MT468698 | ctr pep 31 | GIPCGESCVYIPCTVTALLGCSCKNKVCFRN | 3253.5 | |
| 32 | MT468699 | ctr pep 32 | DTIPCGESCVWIPCISSILGCSCKDKVCYHN | 3348.5 | Cliotide T14 |
| 33 | MT468700/ MT468702 | ctr pep 33 | GIPCGESCVFISCTVTALLGCSCKDKVCYKN | 3216.5 | |
| 34 | MT468703/ MT468704/ MT468709/ MT468724 | ctr pep 34 | DTIPCGESCVFIPCTSSVLGCSCKDKFCYNN | 3308.4 | |
| 35 | MT468705 | ctr pep 35 | GIPCGESCVFIPCTVTALLGCSCKDKVCYKN | 3226.5 | Cter R; CliotideT7 |
| 36 | MT468706 | ctr pep 36 | NIFPCGESCVYIPCITSIVGCSCQNKVCYNN | 3346.5 | |
| 37 | MT468707/ MT468701 | ctr pep 37 | GALCDERCTYVPCISAARGCSCNIHRVCSMN | 3307.4 | Cter 29; CliotideT44 |
| 38 | MT468708 | ctr pep 38 | GGTACGESCIYLPCISGVFEGCSCQNKACYKN | 3280.4 | |
| 39 | MT468710 | ctr pep 39 | GIPCGESCVFIPCTITALLGCSCKDKVCYKN | 3240.5 | Cliotide T11; Cter 21 |
| 40 | MT468711/ MT468712 | ctr pep 40 | SYIPCGESCVYIPCTVTALLGCSCSNKVCYKN | 3393.5 | Cter 10; CliotideT34 |
| 41 | MT468713 | ctr pep 41 | DLICSSTCLHTPCKASVCYCKNAVCYKN | 3042.4 | Cter 18; CliotideT43 |
| 42 | MT468714/ MT468715 | ctr pep 42 | DLQCAETCVHSPCIGPCYCKHGVICYKN | 3059.3 | Cter 31; CliotideT46 |
| 43 | MT468716 | ctr pep 43 | GIPCGESCVFLPCFIIPGCSCKDKVCYLN | 3083.4 | |
| 44 | MT468717 | ctr pep 44 | VDGFCLETCVILPCFSSVAGCYCHGSTCMRG | 3233.4 | Cliotide T37 |
| 45 | MT468720/ MT468719/ MT468718 | ctr pep 45 | GIPCGESCVFIPCISSVVGCSCKSKVCYNN | 3071.4 | Cliotide T8 |
| 46 | MT468721/ MT468726 | ctr pep 46 | GIPCGESCVYIPCTVTALLGCSCKNKVCYRN | 3269.5 | Cliotide T54 |
| 47 | MT468722 | ctr pep 47 | GIPCVESCVFIPCTVTALLGCSCKDKVCYKN | 3268.6 | |
| 48 | MT468723/ MT468725 | ctr pep 48 | GVPCGESCVYIPCTVTALLGCSCKNKVCYRN | 3255.5 | |
| 49 | MT468727/ MT468728/ MT468730 | ctr pep 49 | SIPCGESCVYIPCLTTIVGCSCKSNVCYSN | 3118.4 | Cter 19 |
| 50 | MT468729 | ctr pep 50 | ALPISTIVGCSCKSNVCYSN | 2038.0 | |
| 51 | MT468731 | ctr pep 51 | GVPCAESCVWIPCTVTALLGCSCKDKVCYLN | 3250.5 | Cter B |

**Table S2:** Descriptive statistics of paralysis assay in transgenic *C. elegans* CL4176

| Experiment | Treatment | % of worms not paralyzed at (84th hour) | [a]P value compared to negative control | [b]P value compared positive control |
|---|---|---|---|---|
| 1 | untreated/negative control | 17.42 ± 4.99 | – | <0.001 |
| | vitamin C/positive control | 46.39 ± 7.36 | <0.001 | – |
| 2 | CRF pods (20 µg/ml) | 37.14 ± 10.14 | <0.001 | ns |
| | CRF pods (80 µg/ml) | 47.66 ± 14.86 | <0.001 | ns |
| | CRF pods (200 µg/ml) | 56.21 ± 8.77 | <0.001 | ns |
| 3 | CRF stems (20 µg/ml) | 31.26 ± 5.64 | ns | 0.05 |
| | CRF stems (80 µg/ml) | 43.92 ± 9.52 | <0.001 | ns |
| | CRF stems (200 µg/ml) | 50.70 ± 7.48 | <0.001 | ns |
| 4 | CRF leaves (20 µg/ml) | 28.59 ± 8.30 | ns | 0.006 |
| | CRF leaves (80 µg/ml) | 43.02 ± 8.45 | <0.001 | ns |
| | CRF leaves (200 µg/ml) | 50.57 ± 8.50 | <0.001 | ns |
| 5 | CRF flowers (20 µg/ml) | 31.32 ± 11.52 | ns | 0.05 |
| | CRF flowers (80 µg/ml) | 38.21 ± 10.27 | <0.001 | ns |
| | CRF flowers (200 µg/ml) | 49.62 ± 9.21 | <0.001 | ns |
| 6 | CRF roots (20 µg/ml) | 34.40 ± 10.31 | 0.01 | ns |
| | CRF roots (80 µg/ml) | 40.47 ± 12.51 | <0.001 | ns |
| | CRF roots (200 µg/ml) | 51.30 ± 9.46 | <0.001 | ns |

[a] Statistically significant difference as compared to untreated group
[b] Statistically significant difference as compared to vitamin C treated group

**Table S3:** Descriptive statistics of chemotaxis assay in transgenic *C. elegans* CL2355

| Experiment | Treatment | Chemotaxis index (CI) | [a]P value compared to negative control | [b]P value compared positive control |
|---|---|---|---|---|
| 1 | untreated/negative control | -0.10 ± 0.28 | – | <0.001 |
| | vitamin C/positive control | 0.27 ± 0.13 | <0.001 | – |
| 2 | CRF pods (20 µg/ml) | 0.07 ± 0.14 | ns | ns |
| | CRF pods (80 µg/ml) | 0.16 ± 0.12 | 0.0141 | ns |
| | CRF pods (200 µg/ml) | 0.21 ± 0.12 | 0.0022 | ns |
| 3 | CRF stems (20 µg/ml) | 0.08 ± 0.17 | ns | ns |
| | CRF stems (80 µg/ml) | 0.15 ± 0.11 | 0.0203 | ns |
| | CRF stems (200 µg/ml) | 0.14 ± 0.12 | 0.0297 | 0.0371 |
| 4 | CRF leaves (20 µg/ml) | 0.06 ± 0.11 | ns | ns |
| | CRF leaves (80 µg/ml) | 0.13 ± 0.07 | 0.0305 | ns |
| | CRF leaves (200 µg/ml) | 0.26 ± 0.15 | <0.001 | ns |

| | | | | |
|---|---|---|---|---|
| 5 | CRF flowers (20 µg/ml) | 0.11 ± 0.10 | ns | ns |
| | CRF flowers (80 µg/ml) | 0.15 ± 0.17 | 0.0285 | ns |
| | CRF flowers (200 µg/ml) | 0.17 ± 0.15 | 0.0137 | ns |
| 6 | CRF roots (20 µg/ml) | 0.06 ± 0.20 | ns | ns |
| | CRF roots (80 µg/ml) | 0.13 ± 0.16 | ns | ns |
| | CRF roots (200 µg/ml) | 0.20 ± 0.19 | 0.0121 | ns |

[a] Statistically significant difference as compared to untreated group
[b] Statistically significant difference as compared to vitamin C treated group

**Table S4:** Descriptive statistics of intracellular ROS assay in transgenic *C. elegans* CL2006

| **Experiment** | **Treatment** | **[a]Intracellular ROS (%)** | **[b]P value compared to negative control** | **[c]P value compared positive control** |
|---|---|---|---|---|
| 1 | untreated/negative control | 120.1 ± 5.61 | – | 0.009 |
| | vitamin C/positive control | 107.9 ± 3.5 | 0.009 | – |
| 2 | CRF pods (200 µg/ml) | 104.7 ± 1.02 | <0.001 | ns |
| 3 | CRF stems (200 µg/ml) | 105.4 ± 1.94 | 0.001 | ns |
| 4 | CRF leaves (200 µg/ml) | 103.2 ± 2.03 | <0.001 | ns |
| 5 | CRF flowers (200 µg/ml) | 101.3 ± 6.3 | <0.001 | ns |
| 6 | CRF roots (200 µg/ml) | 102 ± 5.98 | <0.001 | ns |

[a] Percentage of negative control at 0 min
[b] Statistically significant difference as compared to untreated group
[c] Statistically significant difference as compared to vitamin C treated group

**Table S5:** Results of PPCheck webserver for predicting normalized energy per residue for FRODOCK docked models. Best docking pose for each protein-peptide complex is highlighted in green.

**a) 1IYT-2LAM**

| Decoy-ID | Hydrogen Bond Energy (kJ/mol) | Electrostatic Energy (kJ/mol) | van der Waals Energy (kJ/mol) | Total Energy (kJ/mol) | Number of Interface Residues | Normalized Energy per Residue (kJ/mol) |
|---|---|---|---|---|---|---|
| pose2 | 0.00 | -2.58 | -99.83 | -102.41 | 45 | -2.28 |
| pose4 | 0.00 | 0.00 | -101.76 | -101.76 | 49 | -2.08 |
| pose6 | 0.00 | -2.20 | -76.59 | -78.79 | 45 | -1.75 |
| pose5 | 0.00 | 1.06 | -22.65 | -21.58 | 51 | -0.42 |
| pose8 | 0.00 | 0.00 | -20.65 | -20.65 | 52 | -0.40 |
| pose3 | 0.00 | 0.00 | 7.13 | 7.13 | 50 | 0.14 |
| pose10 | 0.00 | 0.00 | 36.74 | 36.74 | 51 | 0.72 |
| pose1 | -8.40 | 0.00 | 86.88 | 78.48 | 46 | 1.71 |
| pose7 | 0.00 | 0.00 | 261.67 | 261.67 | 49 | 5.34 |
| pose9 | 0.00 | 0.00 | 365.36 | 365.36 | 51 | 7.16 |

**b) 2BEG-2LAM**

| Decoy-ID | Hydrogen Bond Energy (kJ/mol) | Electrostatic Energy (kJ/mol) | van der Waals Energy (kJ/mol) | Total Energy (kJ/mol) | Number of Interface Residues | Normalized Energy per Residue (kJ/mol) |
|---|---|---|---|---|---|---|
| pose3 | 0.00 | 0.00 | -91.52 | -91.52 | 49 | -1.87 |
| pose4 | 0.00 | 0.00 | -79.78 | -79.78 | 53 | -1.51 |
| pose10 | 0.00 | 0.00 | -12.82 | -12.82 | 51 | -0.25 |
| pose9 | 0.00 | 0.00 | 74.80 | 74.80 | 61 | 1.23 |
| pose7 | -2.88 | 0.00 | 77.83 | 74.94 | 42 | 1.78 |
| pose5 | 0.00 | 0.00 | 114.80 | 114.80 | 60 | 1.91 |
| pose8 | 0.00 | 0.00 | 174.19 | 174.19 | 43 | 4.05 |
| pose1 | 0.00 | 0.00 | 387.97 | 387.97 | 47 | 8.25 |
| pose2 | -19.84 | 0.00 | 543.28 | 523.44 | 43 | 12.17 |
| pose6 | 0.00 | 0.00 | 819.29 | 819.29 | 67 | 12.23 |

**c) 2MXU-2LAM**

| Decoy-ID | Hydrogen Bond Energy (kJ/mol) | Electrostatic Energy (kJ/mol) | van der Waals Energy (kJ/mol) | Total Energy (kJ/mol) | Number of Interface Residues | Normalized Energy per Residue (kJ/mol) |
|---|---|---|---|---|---|---|
| pose7 | 0.00 | 0.00 | -70.81 | -70.81 | 84 | -0.84 |
| pose6 | -4.13 | 0.00 | -17.44 | -21.57 | 78 | -0.28 |
| pose1 | -16.55 | 0.00 | 139.07 | 122.51 | 89 | 1.38 |
| pose9 | 0.00 | 0.00 | 158.42 | 158.42 | 74 | 2.14 |
| pose8 | 0.00 | 0.00 | 170.22 | 170.22 | 76 | 2.24 |
| pose4 | 0.00 | 0.00 | 302.98 | 302.98 | 74 | 4.09 |
| pose3 | -7.25 | 0.00 | 362.80 | 355.55 | 84 | 4.23 |
| pose10 | 0.00 | 0.00 | 463.90 | 463.90 | 81 | 5.73 |
| pose5 | 0.00 | 0.00 | 616.12 | 616.12 | 94 | 6.55 |
| pose2 | -12.92 | 0.00 | 1034.95 | 1022.03 | 98 | 10.43 |

**d) 2NAO-2LAM**

| Decoy-ID | Hydrogen Bond Energy (kJ/mol) | Electrostatic Energy (kJ/mol) | van der Waals Energy (kJ/mol) | Total Energy (kJ/mol) | Number of Interface Residues | Normalized Energy per Residue (kJ/mol) |
|---|---|---|---|---|---|---|
| pose10 | -4.17 | 0.00 | -95.50 | -99.67 | 65.00 | -1.53 |
| pose5 | -4.78 | 0.00 | -15.15 | -19.94 | 72.00 | -0.28 |
| pose8 | -14.51 | 4.94 | 146.56 | 136.99 | 73.00 | 1.88 |
| pose7 | 0.00 | 0.00 | 135.39 | 135.39 | 68.00 | 1.99 |
| pose1 | 0.00 | 0.00 | 144.70 | 144.70 | 63.00 | 2.30 |
| pose6 | -4.27 | 0.00 | 254.73 | 250.46 | 70.00 | 3.58 |
| pose2 | -18.43 | 12.34 | 337.50 | 331.40 | 84.00 | 3.95 |
| pose4 | 0.00 | 0.00 | 324.05 | 324.05 | 76.00 | 4.26 |
| pose3 | -7.91 | 0.00 | 382.54 | 374.63 | 71.00 | 5.28 |
| pose9 | -8.57 | 0.00 | 1144.61 | 1136.04 | 82.00 | 13.85 |

**e) 2M4J-2LAM**

| Decoy-ID | Hydrogen Bond Energy (kJ/mol) | Electrostatic Energy (kJ/mol) | van der Waals Energy (kJ/mol) | Total Energy (kJ/mol) | Number of Interface Residues | Normalized Energy per Residue (kJ/mol) |
|---|---|---|---|---|---|---|
| pose4 | 0.00 | 0.00 | -67.47 | -67.47 | 95.00 | -0.71 |
| pose9 | -9.08 | 0.00 | -10.26 | -19.33 | 83.00 | -0.23 |
| pose2 | -15.64 | 0.00 | 14.70 | -0.94 | 92.00 | -0.01 |
| pose5 | -6.80 | 0.00 | 61.42 | 54.62 | 94.00 | 0.58 |

| | | | | | | |
|---|---|---|---|---|---|---|
| pose7 | -17.90 | 0.00 | 129.68 | 111.79 | 94.00 | 1.19 |
| pose8 | -36.02 | 0.00 | 152.62 | 116.59 | 87.00 | 1.34 |
| pose1 | -16.73 | 0.00 | 208.95 | 192.22 | 97.00 | 1.98 |
| pose3 | 0.00 | 0.00 | 254.71 | 254.71 | 92.00 | 2.77 |
| pose6 | -89.56 | 0.00 | 659.28 | 569.71 | 95.00 | 6.00 |
| pose10 | 0.00 | 0.00 | 987.48 | 987.48 | 96.00 | 10.29 |

**Figure S2:** Plots of RMSD (top left panel), RMSF (top right panel), Radius of gyration (bottom left panel) and number of hydrogen bonds (bottom right panel) for complexes between cyclotide (PDB ID: 2LAM) and (a) A$\beta_{1\text{-}42}$ monomer (PDB ID: 1IYT), (b) A$\beta_{17\text{-}42}$ U-shaped pentamer (PDB ID: 2BEG), (c) A$\beta_{11\text{-}42}$ S-shaped model (PDB ID: 2MXU), (d) A$\beta_{1\text{-}42}$ disease relevant A$\beta_{1\text{-}42}$ fibrils (PDB ID: 2NAO) (e) A$\beta_{1\text{-}40}$ isolated from brain of AD patient (PDB ID: 2M4J), along the simulation period.

**(a) 1IYT-2LAM**

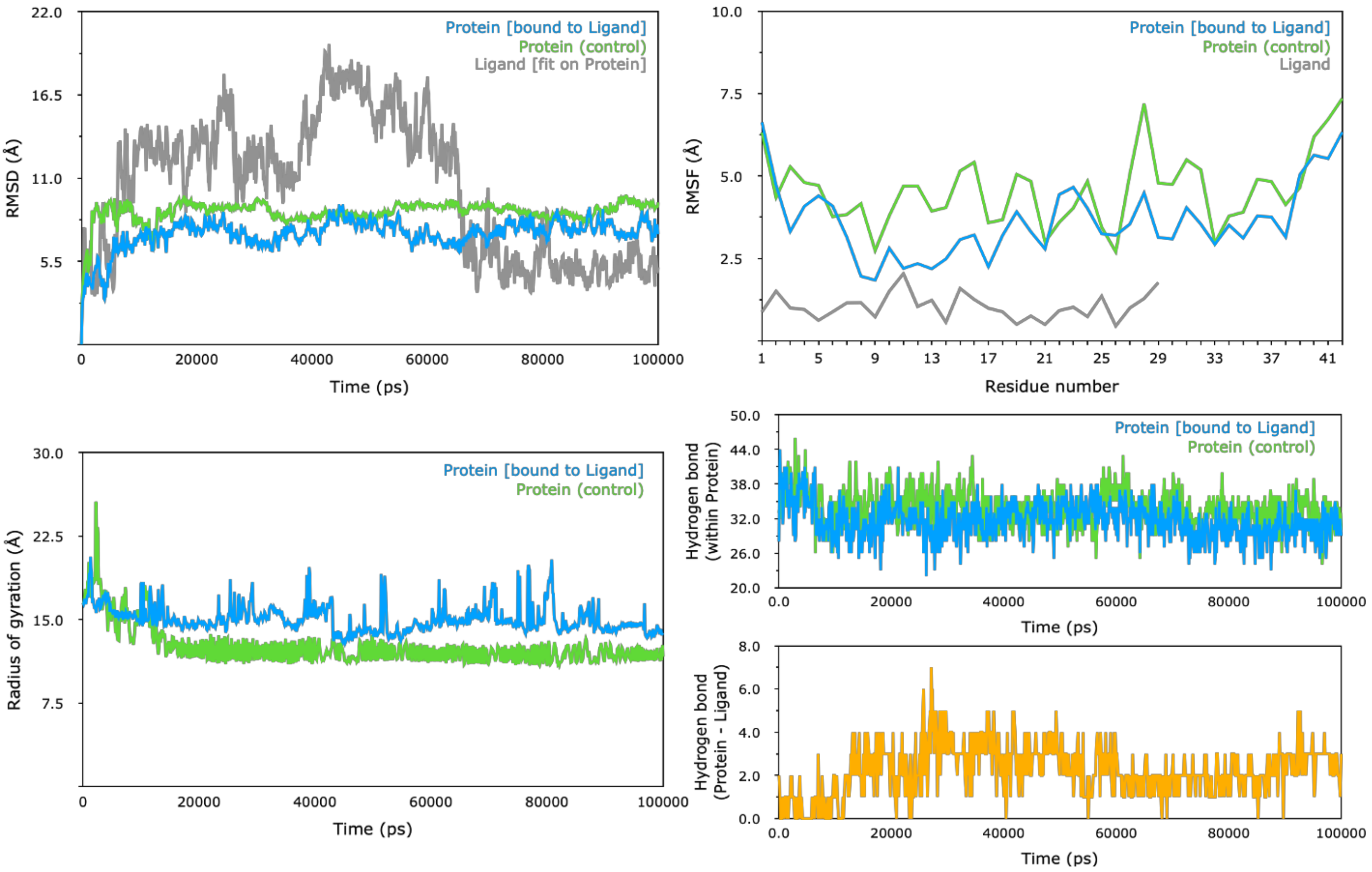

**(b) 2BEG-2LAM**

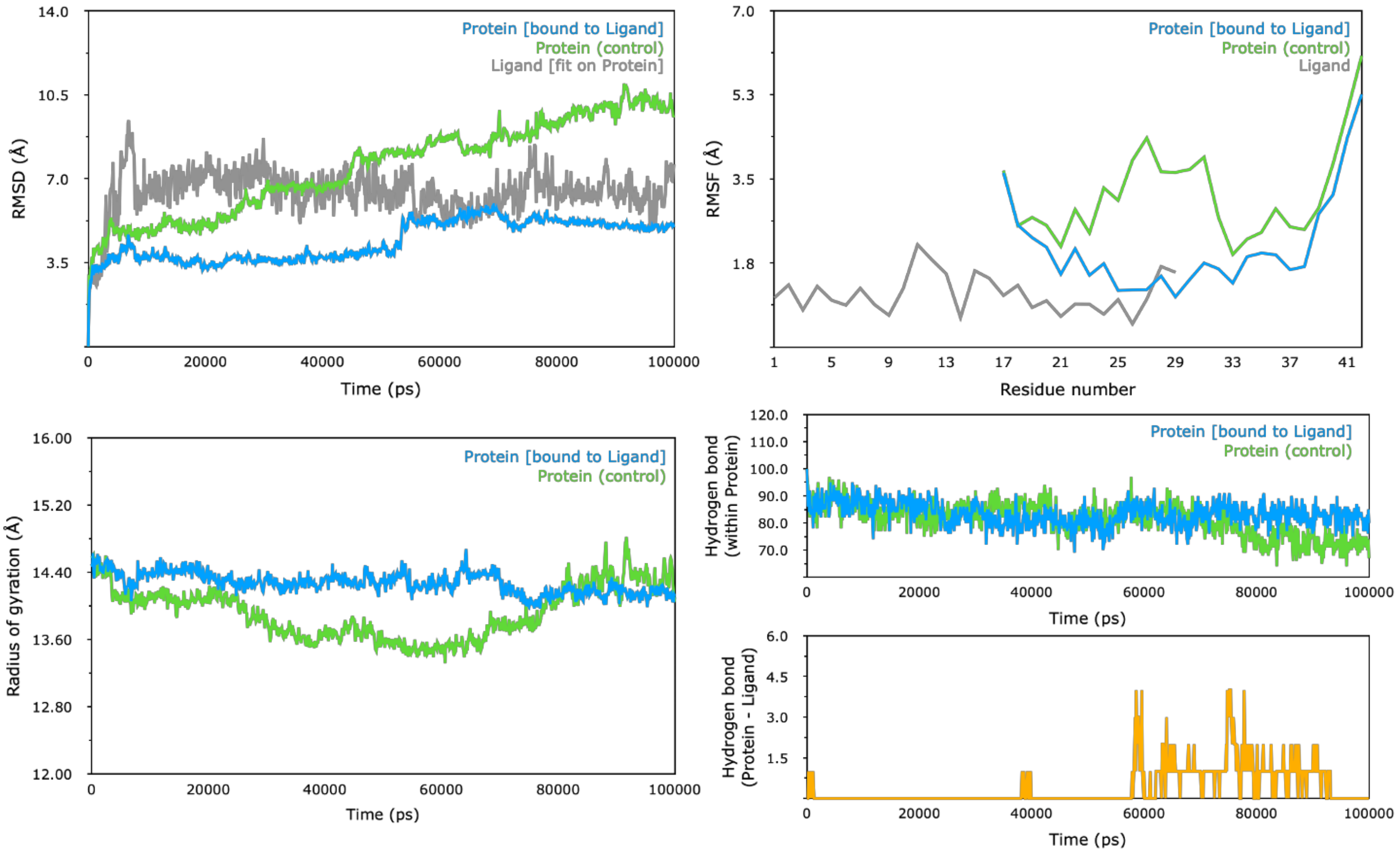


**(c) 2MXU-2LAM**

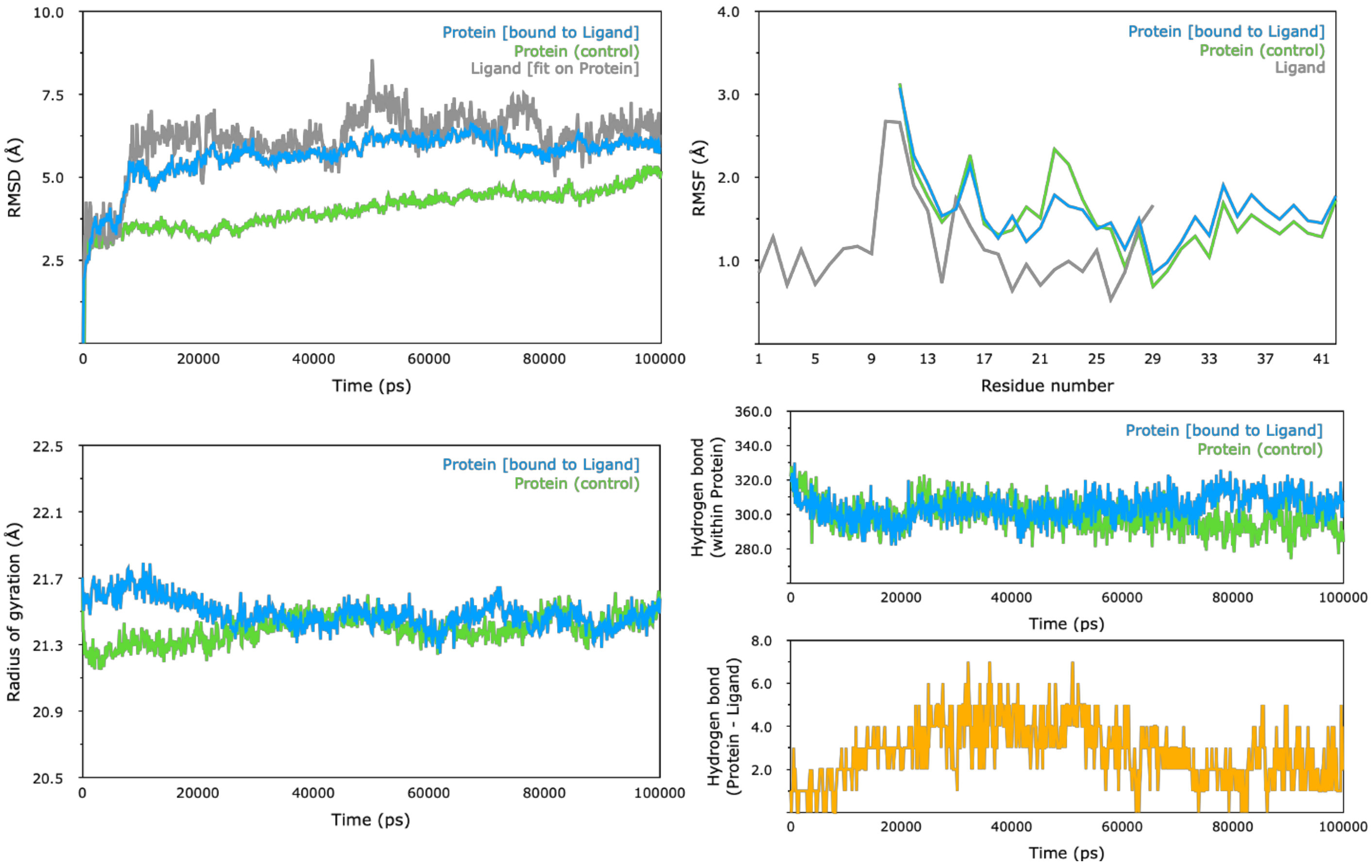

**(d) 2NAO-2LAM**

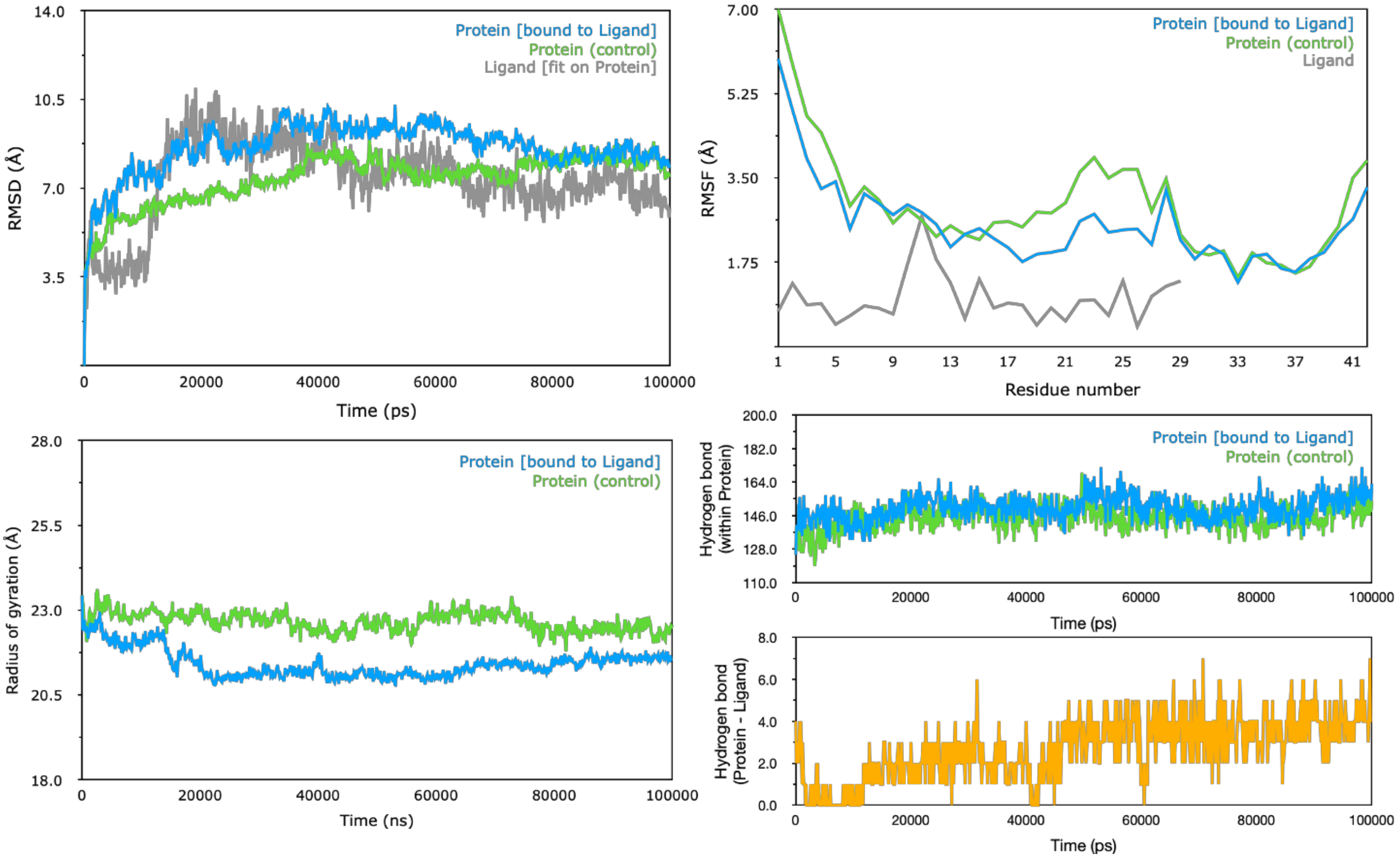


**(e) 2M4J-2LAM**

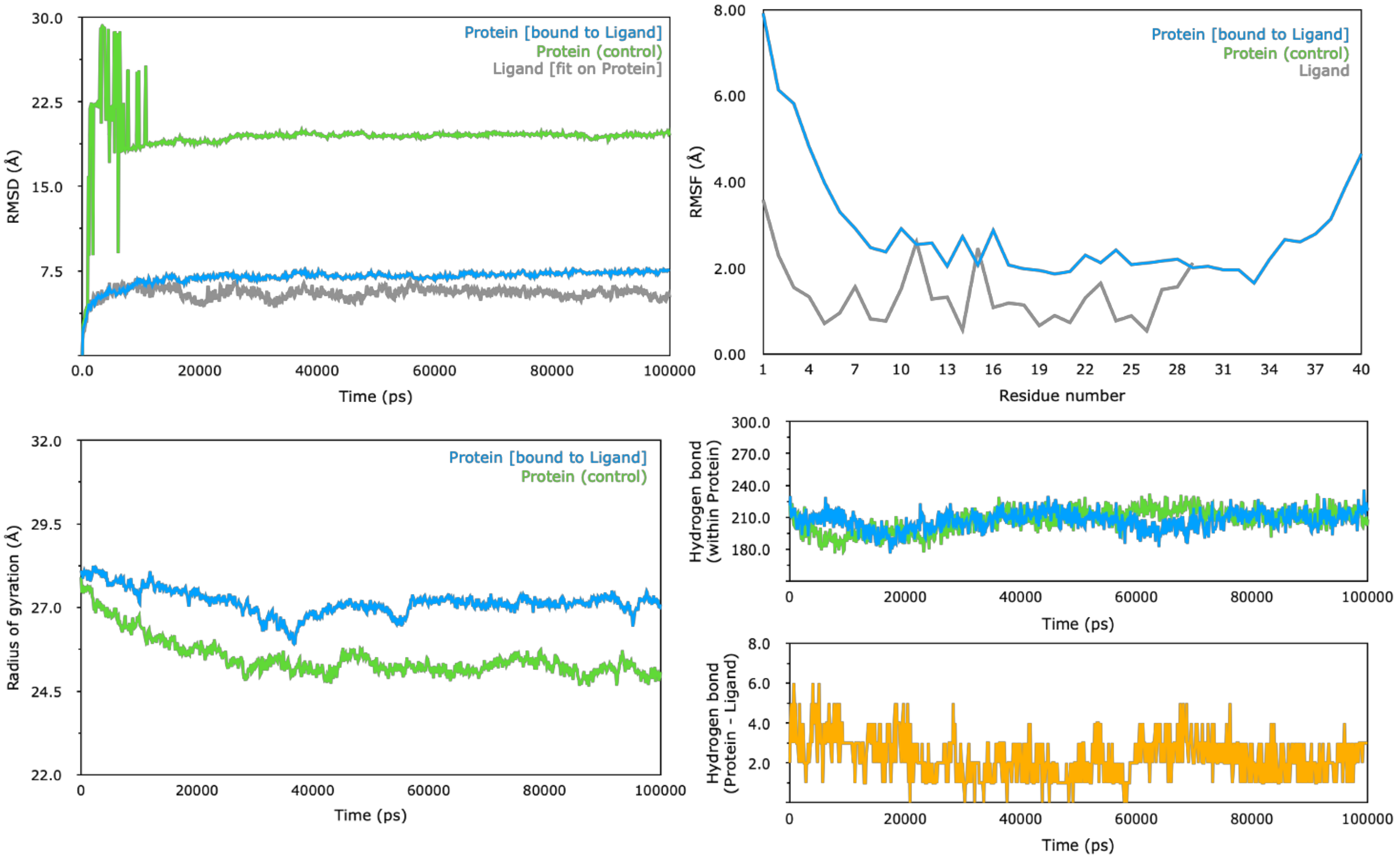

**Figure S3:** Secondary structure timeline analysis during 100 ns MD simulation of (a) A$\beta_{1-42}$ fibril (PDB ID: 2NAO) bound to cyclotide Cter-M (b) unbound A$\beta_{1-42}$ fibril (PDB ID: 2NAO). Colour code explanation of the secondary structures are shown in the small panel below. T denotes turn (aqua); E represents the β-sheet (yellow); B represents isolated bridges (dark yellow); H denotes α-helix (pink); G indicates the $3_{10}$ helix (blue); I denotes π-helix (red) and C indicates random coils (white). These structural analyses were computed using VMD timeline plugin.

**(a)**

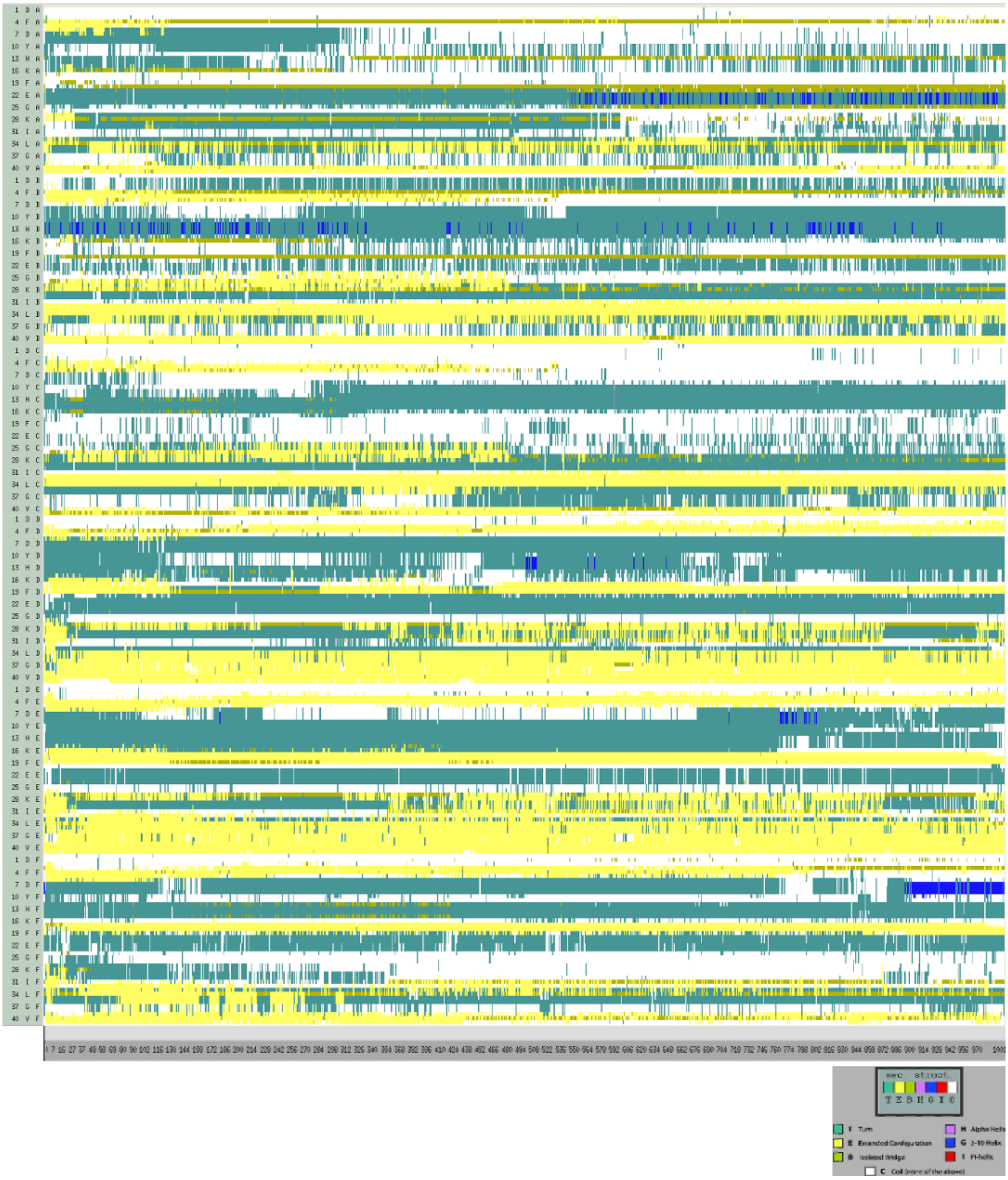

(b)

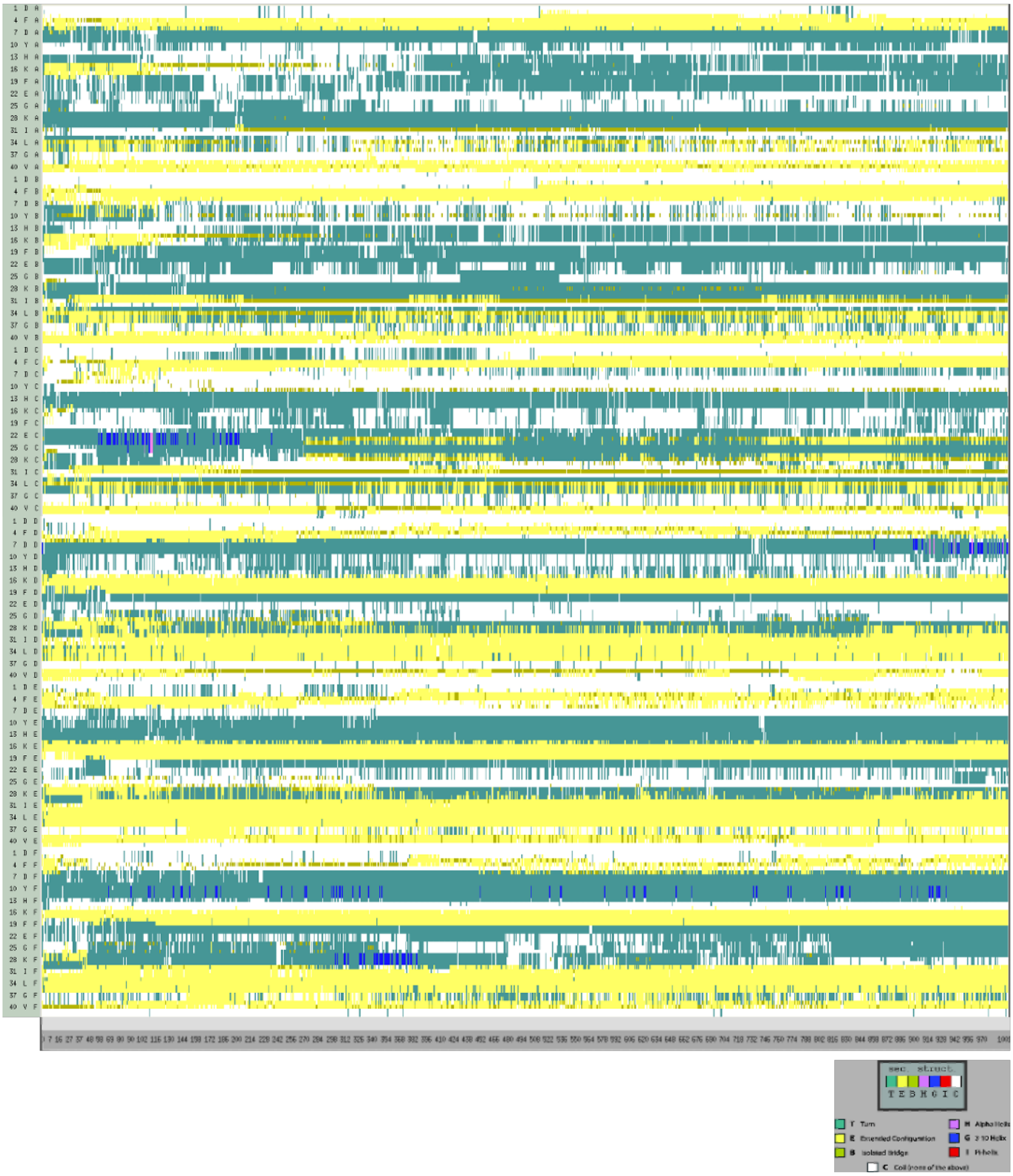
sec. struct.
T E B H G I C
T Turn
E Extended Configuration
B Isolated Bridge
H Alpha Helix
G 3-10 Helix
I Pi-helix
C Coil (none of the above)

**Figure S4:** Snapshots of MD simulation of cyclotide (PDB ID: 2LAM; cyan surface representation) and (a) A$\beta_{1\text{-}42}$ monomer (PDB ID: 1IYT, green cartoon representation), (b) A$\beta_{17\text{-}42}$ U-shaped pentamer (PDB ID: 2BEG, green cartoon representation), (c) A$\beta_{11\text{-}42}$ S-shaped model (PDB ID: 2MXU, green cartoon representation), (d) A$\beta_{1\text{-}40}$ isolated from brain of AD patient (PDB ID: 2M4J, green cartoon representation) complexes at different time points along the simulation period.

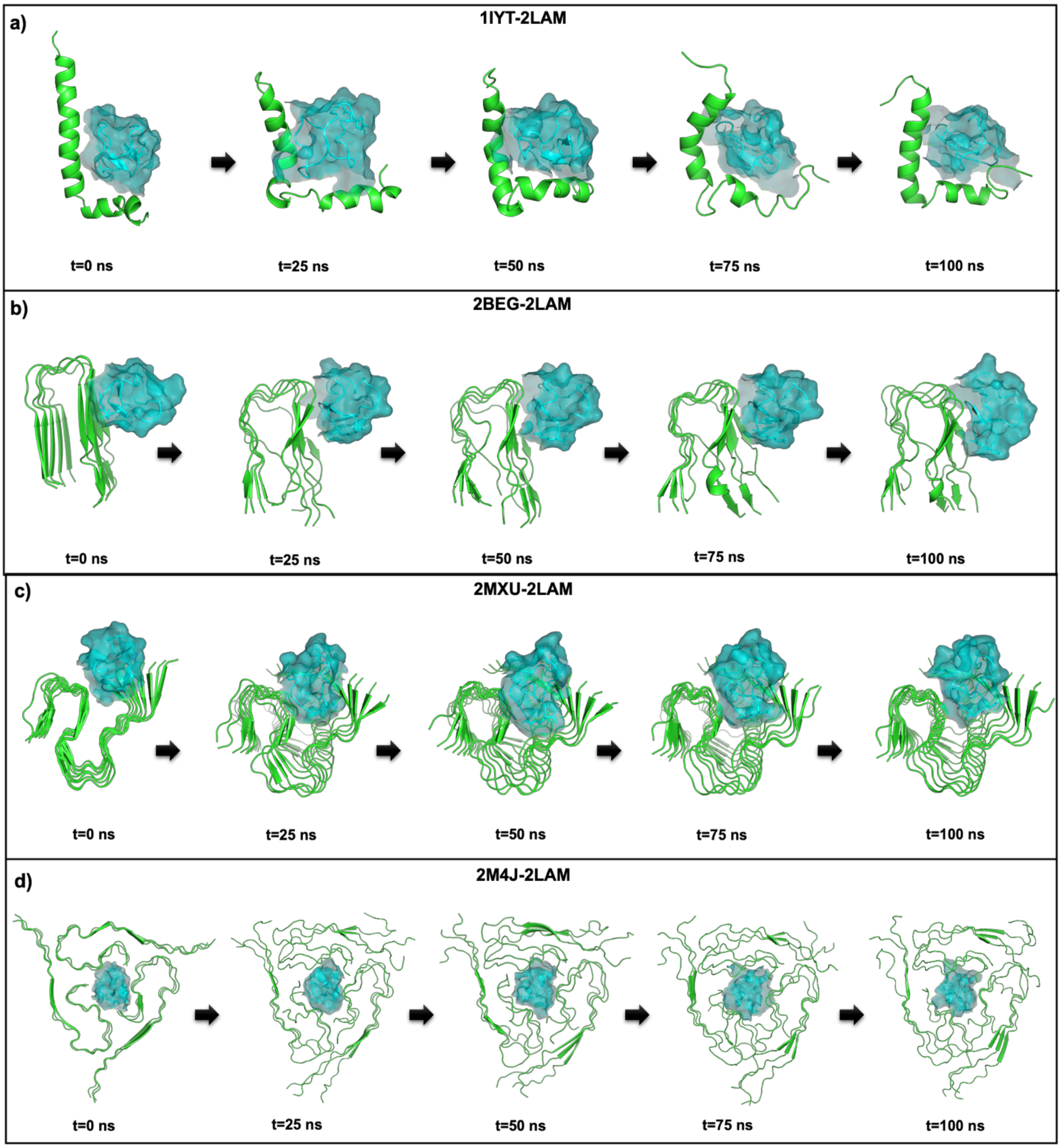

**Table S6:** The total stabilizing energy at t=0 ns and t=100 ns frames for each of the five cyclotide - Aβ complexes.

| Total Stabilizing Energy | 1IYT-2LAM | | 2BEG-2LAM | | 2MXU-2LAM | | 2NAO-2LAM | | 2M4J-2LAM | |
|---|---|---|---|---|---|---|---|---|---|---|
| | t=0 ns | t=100 ns | t=0 ns | t=100 ns | t=0 ns | t=100 ns | t=0 ns | t=100 ns | t=0 ns | t=100 ns |
| Hydrogen Bond Energy (kJ/mol) | 0.00 | 0.00 | 0.00 | 0.00 | -8.83 | -21.18 | -20.12 | -25.95 | -11.17 | -20.33 |
| Electrostatic Energy (kJ/mol) | 0.00 | -21.84 | 0.00 | 0.00 | 0.00 | 0.00 | -4.65 | -6.05 | 0.00 | 0.00 |
| Van der Waals Energy (kJ/mol) | -97.70 | -176.85 | -123.14 | -129.41 | -180.95 | -208.29 | -208.13 | -206.93 | -249.88 | -271.29 |
| Total Stabilizing Energy (kJ/mol) | -97.70 | -198.69 | -123.14 | -129.41 | -189.77 | -229.47 | -232.91 | -238.94 | -261.05 | -291.62 |
| Number of interface residues | 43 | 53 | 52 | 52 | 87 | 91 | 75 | 74 | 101 | 97 |
| Normalized Energy per residue (kJ/mol) | -2.27 | -3.75 | -2.37 | -2.49 | -2.18 | -2.52 | -3.11 | -3.23 | -2.58 | -3.01 |
| No. of Hydrophobic Interactions | 0 | 1 | 2 | 2 | 1 | 4 | 5 | 6 | 3 | 4 |
| No. of van der Waals Pairs | 3791 | 6116 | 5534 | 5736 | 8970 | 8974 | 7768 | 8346 | 11085 | 10841 |
| No. of Salt Bridges | 0 | 1 | 0 | 0 | 0 | 0 | 0 | 0 | 0 | 0 |
| No. of Potential Favourable Electrostatic Interactions | 0 | 2 | 0 | 0 | 0 | 0 | 2 | 3 | 0 | 0 |

**Table S7:** Analysis of protein-protein interactions at t=0 ns (top) and t=100ns (bottom) frames between cyclotide (PDB ID: 2LAM) and $A\beta_{1-42}$ monomer (PDB ID: 1IYT).

| Interactions | [a]Maestro | [a]PPCheck | 1IYT | | | | 2LAM | | | | [b]Type of Bond | Distance Å |
|---|---|---|---|---|---|---|---|---|---|---|---|---|
| | | | Residue-1 | | | | Residue-2 | | | | | |
| | | | Res No. | Res Name | Chain | Atom Name | Res No. | Res Name | Chain | Atom Name | | |
| t=0 ns | | | | | | | | | | | | |
| Hydrogen Bond | ✓ | - | 16 | LYS | A | NZ | 14 | CYS | Z | O | SB | 3.3 |
| | ✓ | - | 16 | LYS | A | NZ | 16 | VAL | Z | O | SB | 2.8 |

| | | | | | | | | | | | |
|---|---|---|---|---|---|---|---|---|---|---|---|
| Aromatic Hydrogen Bond | ✓ | - | 20 | PHE | A | CD2 | 12 | GLY | Z | O | SB | 3.6 |
| | ✓ | - | 20 | PHE | A | CZ | 19 | CYS | Z | O | SB | 3.5 |
| π-π Interactions | ✓ | - | 13 | HIS | H | ring | 15 | TYR | Z | ring | SS | 5.1 |
| **t=100 ns** | | | | | | | | | | | | |
| Hydrogen Bond | ✓ | - | 16 | LYS | A | NZ | 18 | ASP | Z | OD2 | SB | 2.9 |
| | ✓ | - | 13 | HIS | A | ND1 | 15 | TYR | Z | O | SB | 3.0 |
| | ✓ | - | 38 | GLY | A | O | 23 | TRP | Z | NE1 | BS | 2.8 |
| Aromatic Hydrogen Bond | ✓ | - | 13 | HIS | A | CE1 | 14 | CYS | Z | O | SB | 3.4 |
| | ✓ | - | 13 | HIS | A | CE1 | 19 | CYS | Z | O | SB | 3.6 |
| | ✓ | - | 10 | TYR | A | CD1 | 15 | TYR | Z | O | SB | 3.6 |
| | ✓ | - | 40 | VAL | A | O | 23 | TRP | Z | CZ2 | BS | 3.2 |
| Hydrophobic Interactions | - | ✓ | 10 | TYR | A | CB | 15 | TYR | Z | CB | SS | 5.2 |
| Electrostatic Interactions | - | ✓ | 13 | HIS | A | CB | 18 | ASP | Z | CB | SS | 6.6 |
| | - | ✓ | 16 | LYS | A | CB | 18 | ASP | Z | CB | SS | 8.3 |

[a] Protein-protein interaction identified from Maestro and/or PPCheck is highlighted with as ✓.

[b] "SS" represents sidechain-sidechain interaction, "SB" represents sidechain-backbone interaction, and "BB" represents backbone-backbone mode of interaction between the two interacting amino acids.

**Table S8:** PPCheck results for protein-protein interactions at t=0 ns (top) and t=100ns (bottom) frames between cyclotide (PDB ID: 2LAM) and A$\beta_{17\text{-}42}$ U-shaped pentamer (PDB ID: 2BEG).

| **Interactions** | **[a]Maestro** | **[a]PPCheck** | **2BEG** | | | | **2LAM** | | | | **[b]Type of Bond** | **Distance Å** |
|---|---|---|---|---|---|---|---|---|---|---|---|---|
| | | | **Residue-1** | | | | **Residue-2** | | | | | |
| | | | **Res No.** | **Res Name** | **Chain** | **Atom Name** | **Res No.** | **Res Name** | **Chain** | **Atom Name** | | |
| **t=0 ns** | | | | | | | | | | | | |
| Hydrophobic Interactions | - | ✓ | 31 | ILE | C | CB | 25 | ILE | Z | CB | SS | 6.4 |
| | - | ✓ | 31 | ILE | D | CB | 25 | ILE | Z | CB | SS | 6.5 |
| **t=100 ns** | | | | | | | | | | | | |
| Aromatic Hydrogen Bond | ✓ | - | 32 | ILE | B | O | 23 | TRP | Z | CZ3 | BS | 3.3 |
| Hydrophobic Interactions | - | ✓ | 31 | ILE | C | CB | 25 | ILE | Z | CB | SS | 6.6 |
| | - | ✓ | 31 | ILE | D | CB | 25 | ILE | Z | CB | SS | 5.8 |

[a] Protein-protein interaction identified from Maestro and/or PPCheck is highlighted with as ✓.

[b] "SS" represents sidechain-sidechain interaction, "SB" represents sidechain-backbone interaction, and "BB" represents backbone-backbone mode of interaction between the two interacting amino acids.

**Table S9:** PPCheck results for protein-protein interactions at t=0 ns (top) and t=100ns (bottom) frames between cyclotide (PDB ID: 2LAM) and $A\beta_{11-42}$ S-shaped model (PDB ID: 2MXU).

| Interactions | [a]Maestro | [a]PPCheck | 2MXU | | | | 2LAM | | | | [b]Type of Bond | Distance Å |
|---|---|---|---|---|---|---|---|---|---|---|---|---|
| | | | Residue-1 | | | | Residue-2 | | | | | |
| | | | Res No. | Res Name | Chain | Atom Name | Res No. | Res Name | Chain | Atom Name | | |
| **t=0 ns** | | | | | | | | | | | | |
| Hydrogen Bond | ✓ | ✓ | 14 | HIS | K | NE2 | 20 | SER | Z | OG | SS | 3.1 |
| Hydrophobic Interactions | - | ✓ | 34 | LEU | J | CB | 25 | ILE | Z | CB | SS | 5.4 |
| π-π Interactions | ✓ | - | 14 | HIS | H | ring | 23 | TRP | Z | ring 1 | SS | 4.9 |
| | ✓ | - | 14 | HIS | H | ring | 23 | TRP | Z | ring 2 | SS | 4.5 |
| **t=100 ns** | | | | | | | | | | | | |
| Hydrogen Bond | ✓ | - | 11 | GLU | H | N | 12 | GLY | Z | O | BB | 2.8 |
| | - | ✓ | 11 | GLU | H | N | 13 | THR | Z | OG1 | BS | 2.9 |
| | ✓ | ✓ | 14 | HIS | I | NE2 | 21 | CYS | Z | O | SB | 3.1 |
| | - | ✓ | 14 | HIS | J | NE2 | 20 | SER | Z | OG | SS | 3.2 |
| Aromatic Hydrogen Bond | ✓ | - | 19 | PHE | K | CE1 | 1 | GLY | Z | O | SB | 3.9 |
| Hydrophobic Interactions | - | ✓ | 32 | ILE | J | CB | 25 | ILE | Z | CB | SS | 6.8 |
| | - | ✓ | 34 | LEU | J | CB | 25 | ILE | Z | CB | SS | 6.4 |
| | - | ✓ | 34 | LEU | K | CB | 25 | ILE | Z | CB | SS | 5.2 |
| π-π Interactions | ✓ | - | 14 | HIS | G | ring | 23 | TRP | Z | ring 2 | SS | 5.2 |

[a] Protein-protein interaction identified from Maestro and/or PPCheck is highlighted with as ✓.

[b] "SS" represents sidechain-sidechain interaction, "SB" represents sidechain-backbone interaction, and "BB" represents backbone-backbone mode of interaction between the two interacting amino acids.

**Table S10:** PPCheck results for protein-protein interactions at t=0 ns (top) and t=100ns (bottom) frames between cyclotide (PDB ID: 2LAM) and $A\beta_{1-40}$ isolated from brain of AD patient (PDB ID: 2M4J).

| Interactions | [a]Maestro | [a]PPCheck | 2M4J Residue-1 | | | | 2LAM Residue-2 | | | | [b]Type of Bond | Distance Å |
|---|---|---|---|---|---|---|---|---|---|---|---|---|
| | | | Res No. | Res Name | Chain | Atom Name | Res No. | Res Name | Chain | Atom Name | | |
| **t=0 ns** | | | | | | | | | | | | |
| Hydrogen Bond | ✓ | ✓ | 40 | VAL | E | OXT | 18 | ASP | Z | N | SB | 2.7 |
| | ✓ | - | 40 | VAL | D | O | 1 | GLY | Z | N | BB | 2.7 |
| Hydrophobic Interactions | - | ✓ | 32 | ILE | B | CB | 16 | VAL | Z | CB | SS | 5.5 |
| | - | ✓ | 32 | ILE | C | CB | 25 | ILE | Z | CB | SS | 6.2 |
| | - | ✓ | 32 | ILE | I | CB | 2 | LEU | Z | CB | SS | 6.3 |
| Salt Bridges | ✓ | - | 40 | VAL | D | OXT | 1 | GLY | Z | N | SB | 3.2 |
| | ✓ | - | 40 | VAL | G | OXT | 1 | GLY | Z | N | SB | 4.3 |
| **t=100 ns** | | | | | | | | | | | | |
| Hydrogen Bond | ✓ | ✓ | 35 | MET | B | N | 15 | TYR | Z | O | BB | 2.8 |
| | ✓ | ✓ | 32 | ILE | D | O | 21 | CYS | Z | N | BB | 2.8 |
| | ✓ | - | 33 | GLY | I | N | 29 | ASN | Z | OD1 | SB | 3.4 |
| Aromatic Hydrogen Bond | ✓ | - | 32 | ILE | A | O | 23 | TRP | Z | ring | SB | 4.2 |
| Hydrophobic Interactions | - | ✓ | 32 | ILE | B | CB | 16 | VAL | Z | CB | SS | 6.4 |
| | - | ✓ | 32 | ILE | C | CB | 25 | ILE | Z | CB | SS | 5.3 |
| | - | ✓ | 32 | ILE | F | CB | 25 | ILE | Z | CB | SS | 5.3 |
| | - | ✓ | 39 | VAL | F | CB | 2 | LEU | Z | CB | SS | 5.9 |

[a] Protein-protein interaction identified from Maestro and/or PPCheck is highlighted with as ✓.

[b] "SS" represents sidechain-sidechain interaction, "SB" represents sidechain-backbone interaction, and "BB" represents backbone-backbone mode of interaction between the two interacting amino acids.

**Figure S5:** Molecular interactions between cyclotide (PDB ID: 2LAM; chain Z; cyan cartoon representation) and (a) A$\beta_{1\text{-}42}$ monomer (PDB ID: 1IYT; chain A; green cartoon representation), (b) A$\beta_{17\text{-}42}$ U-shaped pentamer (PDB ID: 2BEG; chain A-E; green cartoon representation), (c) A$\beta_{11\text{-}42}$ S-shaped model (PDB ID: 2MXU; chain A-L; green cartoon representation), (d) A$\beta_{1\text{-}40}$ isolated from brain of AD patient (PDB ID: 2M4J; chain A-I; green cartoon representation) at the beginning ($0^{th}$ ns snapshot) and end ($100^{th}$ ns snapshot) of the simulation period. Colour scheme for interactions used: classical hydrogen bonds in black, aromatic hydrogen bonds in green, hydrophobic interactions in blue, electrostatic interactions in orange, salt-bridges in pink and π-π interactions in red.

**a) 1IYT (A$\beta_{1\text{-}42}$ monomer) - 2LAM (cyclotide) complex**

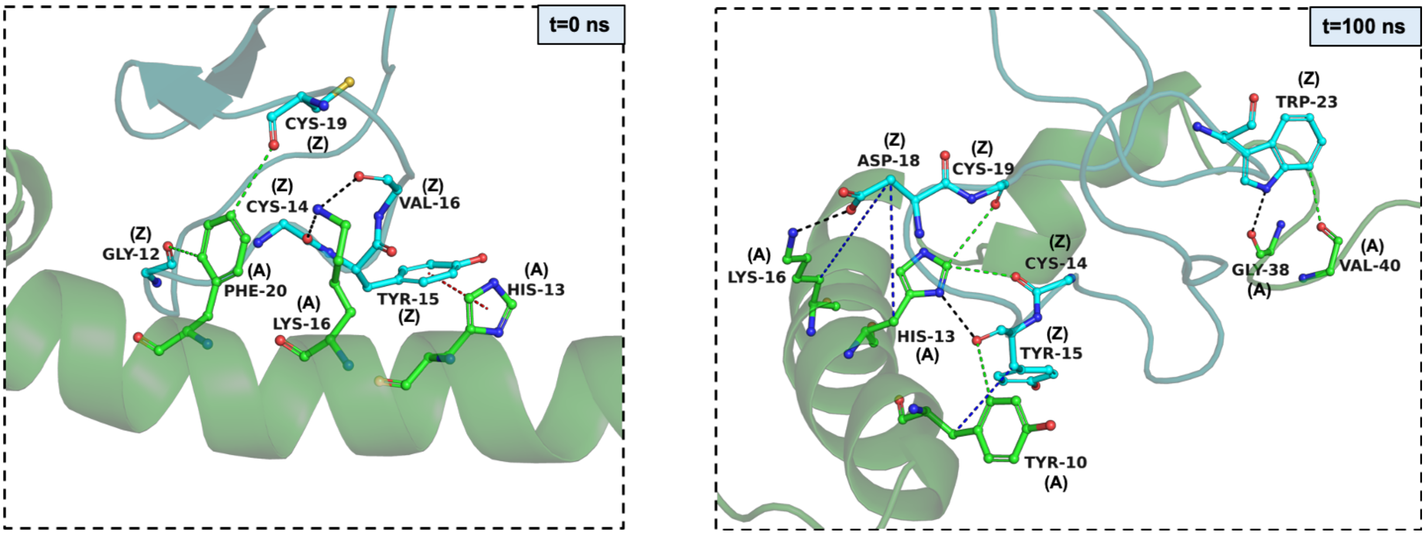


**b) 2BEG (A$\beta_{17\text{-}42}$ fibril) - 2LAM (cyclotide) complex**

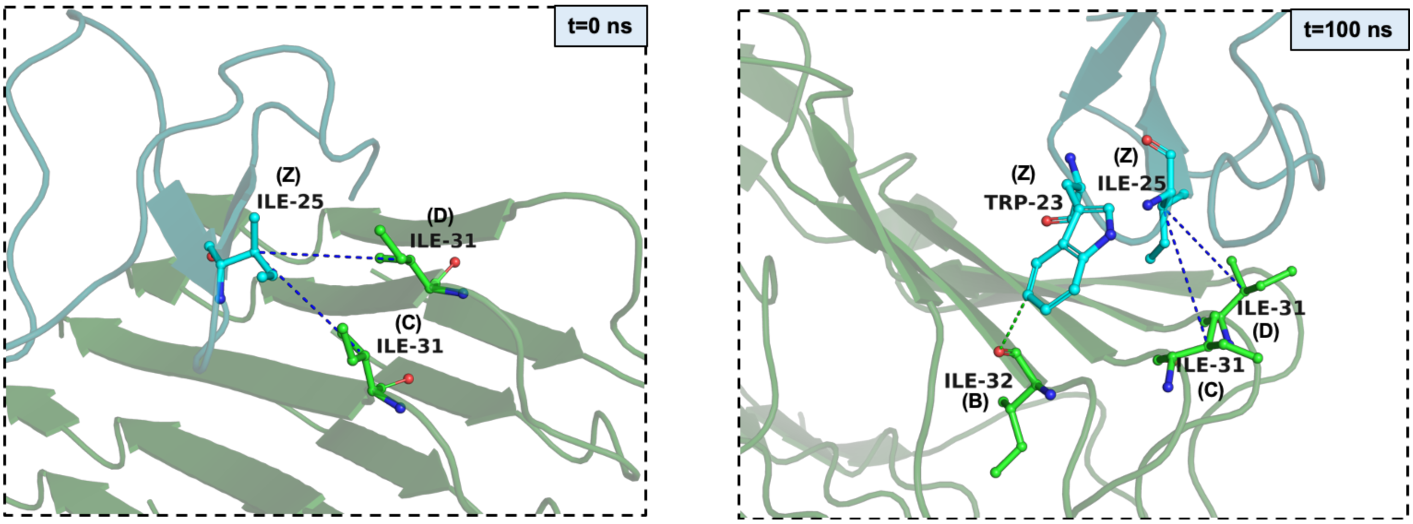

c) 2MXU (A$\beta_{11-42}$ fibril) - 2LAM (cyclotide) complex

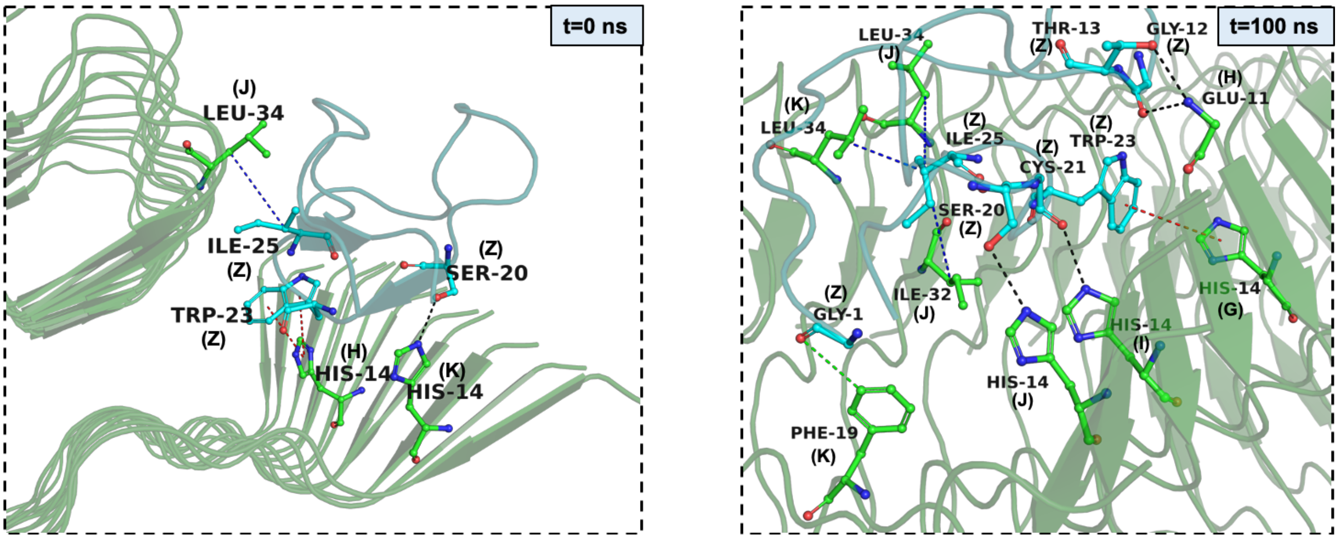


d) 2M4J (A$\beta_{1-40}$ fibril) - 2LAM (cyclotide) complex

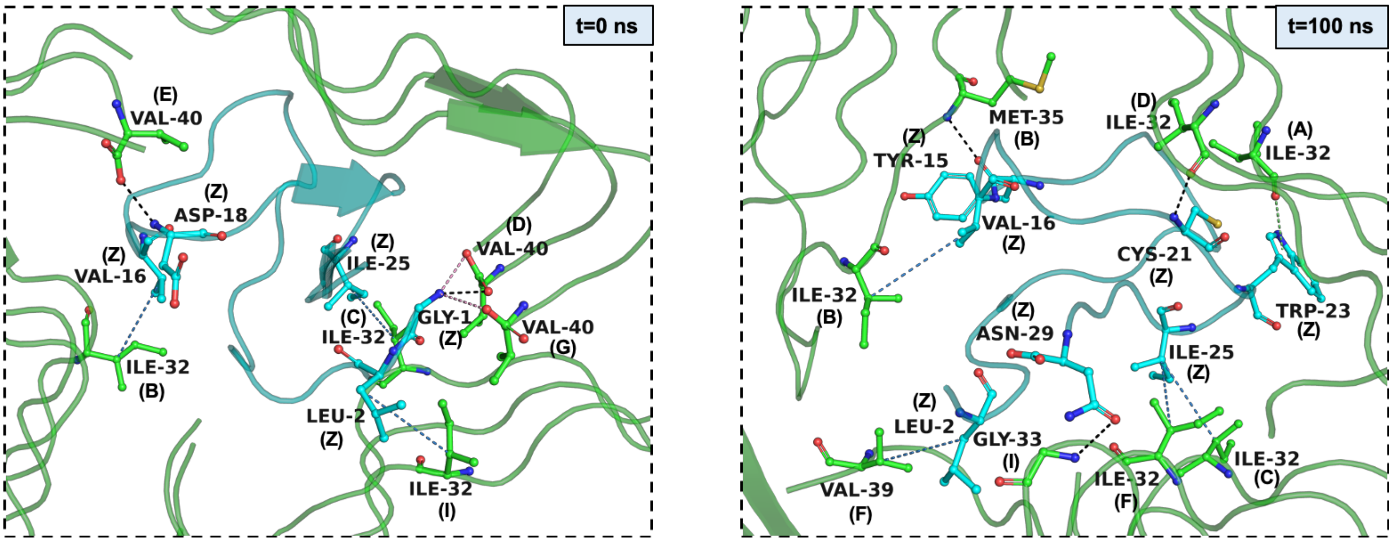
